# eIF2B Activator Rescues Neonatal Lethality of an eIF2Bα Sugar Phosphate Binding Mutation Associated with Vanishing White Matter Disease

**DOI:** 10.1101/2023.05.06.539602

**Authors:** James J. Lee, Nina Ly, Rejani B. Kunjamma, Holly M. Robb, Eric G. Mohler, Janani Sridar, Qi Hao, José Zavala-Solorio, Chunlian Zhang, Ganesh Kolumam, Nick van Bruggen, Caitlin F. Connelly, Carmela Sidrauski

## Abstract

eIF2B is a decameric guanine nucleotide exchange factor (GEF) that is essential for protein synthesis and a key effector of the integrated stress response (ISR). Hypomorphic mutations in any of the eIF2B subunits are associated with Vanishing White Matter Disease (VWM), a leukodystrophy characterized by ISR activation and white matter loss. Here, we showed that the VWM-associated N208Y eIF2Bα mutation, which abolishes sugar phosphate binding, led to a drastic reduction in its level in cells and concomitant ISR activation. We found that N208Y homozygous mice are small and die shortly after birth. Remarkably, continuous availability of 2BAct, a small molecule eIF2B activator, in food rescued the lethality and significantly extended their lifespan. 2BAct-maintained N208Y homozygous mice, however, developed motor deficits and loss of myelin with age. As is the case for milder VWM models, ISR induction was restricted to the central nervous system in treated animals. Upon 2BAct withdrawal, adult mutant mice deteriorated quickly, the ISR was induced in all peripheral tissues tested and resulted in high levels of circulating FGF21 and GDF15. This model provides a novel platform to study the impact of ISR activation across tissues with temporal control.

## Introduction

Eukaryotic cells encounter a variety of insults that lead to activation of the integrated stress response (ISR), a conserved signaling pathway that decreases the protein synthesis rate and induces a broad transcriptional response. Four kinases with distinct sensor domains, PERK, HRI, GCN2 and PKR, are activated by diverse inputs and converge in the phosphorylation of serine 51 in the α subunit of eukaryotic initiation factor 2 (eIF2), a GTPase that is responsible for bringing the initiator met-tRNAi to the ribosome for mRNA translation initiation. This phosphorylation event converts eIF2α from a substrate to an inhibitor of its dedicated guanine nucleotide exchange factor (GEF) eIF2B. As a result, the enzymatic activity of eIF2B is reduced, leading to a decrease in eIF2-GTP-met-tRNAi ternary complex and, consequently, mRNA translation (Algire et al., 2005; Krishnamoorthy et al., 2001). Concomitant with a reduction in bulk protein synthesis upon stress, there is a paradoxical increase in translation of a subset of transcripts containing unique 5’UTRs, which include the transcription factor ATF4 that triggers transcriptional upregulation of ISR target genes (Harding et al., 2000).

The eIF2B complex, a key translation initiation factor and effector of the ISR, is a dimer of pentamers with a central hexameric regulatory core (α_2_β_2_δ_2_) composed of three different subunits that has homology to an archeal bacteria metabolic enzyme. The hexameric core is decorated by two catalytic sub-complexes (γε) and contains the largest subunit, ε, which includes the catalytic domain (Tsai et al., 2018; Zyryanova et al., 2018). Mutations in any of the five eIF2B subunits are associated with Vanishing White Matter Disease (VWM), an ultra rare inherited leukodystrophy characterized by the progressive disappearance of white matter (Hamilton et al., 2018; Shimada et al., 2015). Early onset disease (early childhood) is the most common and is fast progressing and fatal, whereas later onset disease (adolescence and adulthood) is milder and progresses more slowly (Hamilton et al., 2018). More than 60% of patients are compound heterozygotes for mutations in the same subunit of eIF2B and disease progression is quite variable and likely scales with the levels of residual eIF2B activity (Slynko et al., 2021).

Biochemical and cell biological studies revealed that VWM mutations reduce eIF2B complex stability and GEF enzymatic activity (Wong et al., 2018). Homozygous (HOM) mice carrying three different VWM mutations have been generated and characterized and include R484W eIF2Bδ, R191H eIF2Bε and R132H eIF2Bε (Dooves et al., 2016; Geva et al., 2010; Wong et al., 2019). R484W eIF2Bδ HOM and R191H eIF2Bε HOM mice develop VWM pathology, including progressive motor deficit and demyelination with age, and show chronic activation of the ISR that is restricted to the central nervous system (CNS) and primarily in astrocytes and oligodendrocytes (Dooves et al., 2016; Wong et al., 2019). By contrast, R132H eIF2Bε HOM mice have a very mild ISR in the brain and show a developmental delay but do not develop VWM pathology (Geva et al., 2010; Wong et al., 2019). These VWM-associated hypomorphic mutations decrease the stability and GEF activity of the eIF2B complex in cells and lead to chronic activation of the ISR. Importantly, the negative feedback loop of the ISR, which is elicited through activation of the eIF2α phosphatase GADD34, is not effective in turning the pathway off. In this neurodevelopmental disease, the genetic defect that leads to a reduction in eIF2-GTP-met-tRNAi ternary complex formation and ISR activation is downstream of stress-induced eIF2α phosphorylation.

ISRIB-like molecules (including 2BAct) bind to the central core of the regulatory sub-complex bridging the two halves of the eIF2B decamer together, stabilizing the full complex and boosting its GEF activity (Kashiwagi et al., 2019; Kenner et al., 2019; Sidrauski et al., 2013; Sidrauski et al., 2015). We previously showed that preventive treatment of R191H eIF2Bε HOM mice (hereafter referred to as R191H^HOM^) with an orally bioavailable eIF2B activator, 2BAct, prevents development of VWM pathophysiology, fully suppresses the ISR in the brain, and normalizes both the transcriptome and proteome (Wong et al., 2019). We also showed that ISRIB and 2BAct were able to boost the GEF activity of a purified human R195H eIF2Bε (R191H in mice) recombinant complex and of lysates derived from R191H^HOM^ mouse embryonic fibroblasts to wildtype levels (Wong et al., 2019; Wong et al., 2018). The full rescue of the behavioral phenotype in R191H^HOM^ mice by 2BAct underscored the remarkable ability of this mechanism of action to restore eIF2B function, as the corresponding mutation in humans (R195H eIF2Bε) is characterized by a rapidly fatal and early onset disease (Cree leukoencephalopathy).

The α regulatory subunit of eIF2B plays a key role in the assembly of the full complex (Schoof et al., 2021). It forms a homodimer (α_2_) *in vitro* and in cells and bridges the two halves of the eIF2B complex promoting decamer formation, which is the most active form of the enzyme (Wong et al., 2018; Wortham et al., 2014). We recently discovered that, as suggested by its homology to an archeal metabolic enzyme, sugar phosphates bind to eIF2Bα and further boost the enzymatic activity of the complex (Hao et al., 2021). Similar to ISRIB, binding of sugar phosphates stabilizes the eIF2B complex but the synthetic ligand and metabolite exert their action through different binding sites. The impact of sugar phosphate-stimulated eIF2B GEF activity on normal physiology is not known but suggests coupling of activation of the ISR to the metabolic state of cells and in particular, the accumulation and depletion of glycolytic intermediates. A single amino acid substitution in the sugar phosphate binding pocket of the α subunit (N208Y) abolished binding and boosting of the GEF activity by sugar phosphates (Hao et al., 2021). Interestingly, in humans, the N208Y mutation (622A>T) in the *EIF2B1* locus has been reported as a compound heterozygous with a second mutation in the same locus in a single VWM patient but, unfortunately, disease severity of this patient was not reported (van der Knaap et al., 2002).

Here, we report the cellular and organismal effects of a sugar phosphate binding and VWM-associated mutation (N208Y in eIF2Bα). We demonstrate that the ability to bind sugar phosphates is required to maintain normal levels of eIF2Bα in cells. When the N208Y mutation was introduced into mice and bred to homozygosity, it resulted in neonatal lethality. Remarkably, treatment with 2BAct rescued the early lethality and provided life extension for mutant mice beyond 8 months. This is a novel and severe model of VWM disease, akin to early lethality alleles in humans, and we show that 2BAct can boost the GEF activity of N208Y eIF2Bα mutant lysates, although not to normal levels. The inability to fully rescue normal GEF activity results in chronic activation of the ISR only in the CNS and loss of white matter with age in 2BAct-maintained N208Y HOM mice, thus recapitulating the phenotype of milder mouse models of disease. Removal of the drug resulted in marked ISR induction across all tissues (central and peripheral) and led to quick physical deterioration, underscoring the importance of sustained treatment for proper physiological function. This new mouse model will enable the study of the impact of activation of the ISR across cell types and tissues, both central and peripheral, with temporal control by manipulating drug availability.

## Results

### Mutations that reduce sugar phosphate binding lead to a decrease in the level of eIF2Bα and induction of the ISR

To determine the impact of the VWM-associated N208Y mutation in eIF2Bα on eIF2B complex stability and activity in mammalian cells, we introduced this allele into the MIN-6 mouse β-cell line using CRISPR/Cas9. In addition, we introduced a previously characterized mutation E198K that, in contrast to N208Y, only partially impairs fructose-6-phosphate (F6P) binding to eIF2Bα (Hao et al., 2021). We were able to generate two independent clones for each sugar phosphate binding mutation; importantly, supplementation of the media with the eIF2B activator ISRIB during mutagenesis and clonal selection was required. The homozygosity of the point mutations were confirmed by Sanger sequencing (Figure 1A).

**Figure 1.**
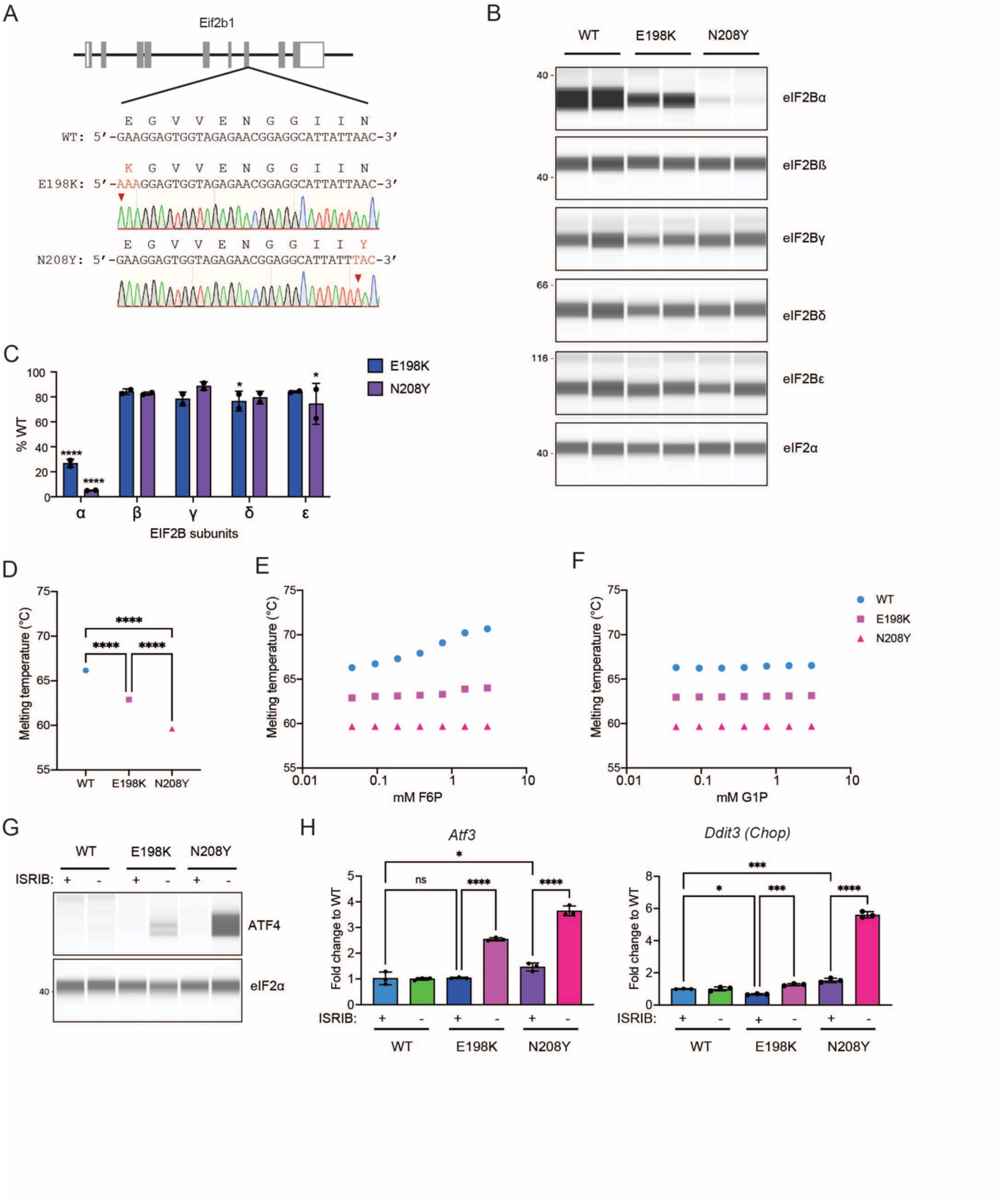
Mutations that reduce sugar phosphate binding lead to a decrease in the level of eIF2Bα and an induction of the ISR. (A) Schematic representation of mouse *Eif2b1* locus and nucleotide substitutions to generate E198K or N208Y mutated MIN-6 cell lines. Solid gray bars represent open reading frames, open bars represent UTRs. A representative sequencing result of a homozygous clone for each mutation is shown. (B) Immunoblot analysis of eIF2B subunits of MIN-6 WT, E198K or N208Y cells maintained in 200 nM ISRIB. Data shown are representative of two clones per genotype. (C) Quantification of the eIF2B subunit level shown in (B). eIF2B subunit expression was normalized to eIF2ɑ and expressed as % of WT. Note reduced level of eIF2Bα in the mutants as compared to wild-type cells. Significance is shown for comparisons of each mutant to WT. n=2. Error bars are SD. Two-way ANOVA with Holm-Sidak’s multiple comparisons test ****p<0.0001, *p<0.05. (D) Calculated midpoint thermal unfolding of recombinant human WT, E198K or N208Y eIF2Bα using NanoDSF without sugar phosphate and with F6P (E) or G1P (F). n=3. Error bars (unable to be displayed) are standard deviation. Standard deviation: WT (0.0001), E198K (0.0002), N208Y (0.0006). One-way ANOVA with Tukey’s multiple comparisons test. **** p < 0.0001 (G) Immunoblot of ATF4 protein and eIF2a before and after 24 hr ISRIB withdrawal. (H) Transcript levels of ISR target genes, *Atf3* and *Ddit3*, after 24 hr ISRIB withdrawal. n=3 of one representative clone. Error bars are SD. One-way ANOVA with Tukey’s multiple comparisons test. * p < 0.05, *** p < 0.001, **** p < 0.0001.

Given that the generation of a yeast strain carrying the corresponding VWM mutation (N209Y) had been reported to significantly reduce the levels of eIF2Bα (Richardson et al., 2004), we first measured the abundance of each eIF2B subunit in the two mutant cell lines. We observed a drastic reduction in the level of eIF2Bα in both sugar phosphate binding mutants, whereas the level of other complex subunits was only slightly affected as compared to wild type (WT) cells (Figure 1B and 1C). The magnitude of the reduction in the level of eIF2Bα in cells correlated with the decrease in the ability of each mutant eIF2B complex to bind F6P *in vitro* (Hao et al., 2021). Whereas N208Y abolished F6P binding and resulted in a 95% decrease in the level of eIF2Bα, E198K retained the ability to bind sugar phosphates, although with a 10-fold decrease in affinity as compared to WT (WT + F6P: Kd = 9.4 ± 2.3 µM and E198K + F6P : Kd = 99 ± 12 µM)(Hao et al., 2021), and resulted in a 74% reduction in subunit abundance (Figure 1C). Western blot of purified N208Y and E198K eIF2B⍺ dimers confirmed that the sugar phosphate binding mutations did not disrupt the ability of the antibody to recognize the epitope (Figure Supplement 1A).

The reduction of eIF2B⍺ in mutant cells could be due to an inherent instability of the sugar phosphate binding mutant proteins. To measure protein stability, we performed nano differential scanning fluorimetry (NanoDSF) of purified recombinant WT, N208Y and E198K eIF2B⍺. This technique measures the intrinsic fluorescence of tryptophan as the protein unfolds as a function of temperature to calculate the midpoint of thermal unfolding (Tm). In the absence of F6P, purified WT eIF2B⍺ protein had a melting temperature of 66.2°C and both sugar phosphate binding mutants showed a significant reduction in thermal stability (E198K: 62.9°C and N208Y: 59.6°C, Figure 1D). As we previously reported (Hao et al., 2021), addition of F6P further increased the stability of WT. However, it did not enhance the stability of either E198K or N208Y eIF2Bα (Figure 1E). Glucose-1-phosphate, which we previously showed did not bind to or boost the activity of eIF2B, did not increase the Tm of any of the purified proteins (Figure 1F). Taken together, these results suggest that mutations that disrupt the sugar phosphate binding pocket, and in particular VWM-associated N208Y, have a significant effect on the intrinsic stability of eIF2Bα and impair further stabilization by metabolite binding.

As all eIF2B subunits are essential in mammalian cells, the striking reduction in eIF2B⍺ levels observed in N208Y MIN-6 cells explained the need to supplement the media with ISRIB during their generation. To further investigate the effect of ISRIB, we measured the status of ISR activation before and after drug withdrawal from the supplemented media. As seen in Figure 1G, we did not detect ATF4 protein in cells grown in the presence of this eIF2B activator in any genotype. However, there was a large increase in ATF4 expression in N208Y cells and to a lesser extent in E198K cells when ISRIB was withdrawn from the media. The upregulation of the ISR resulted in transcriptional induction of *Atf3* in both sugar phosphate binding mutants and the pro-apoptotic gene *Chop* only in N208Y cells (Figure 1H). The significant upregulation of the ISR in N208Y MIN-6 cells and requirement of ISRIB for their generation suggests that this is a particularly severe VWM-associated pathogenic allele.

### Neonatal lethality is observed in N208Y homozygous mice

To determine the impact of sugar phosphate binding *in vivo*, we generated a N208Y mutant mouse strain via targeted mutagenesis (Figure 2A). Heterozygous *Eif2b1^N208Y/+^* (hereafter referred to as N208Y^HET^) mice were viable, grew and bred normally. Breeding of N208Y^HET^ males and females yielded pups of all three genotypes at the expected mendelian ratio at birth but no viable homozygous *Eif2b1^N208Y/N208Y^*(hereafter referred to as N208Y^HOM^) pups at weaning (Figure 2B). Daily monitoring and genotyping of the pups revealed that all N208Y^HOM^ pups died within 24 hours after birth.

**Figure 2.**
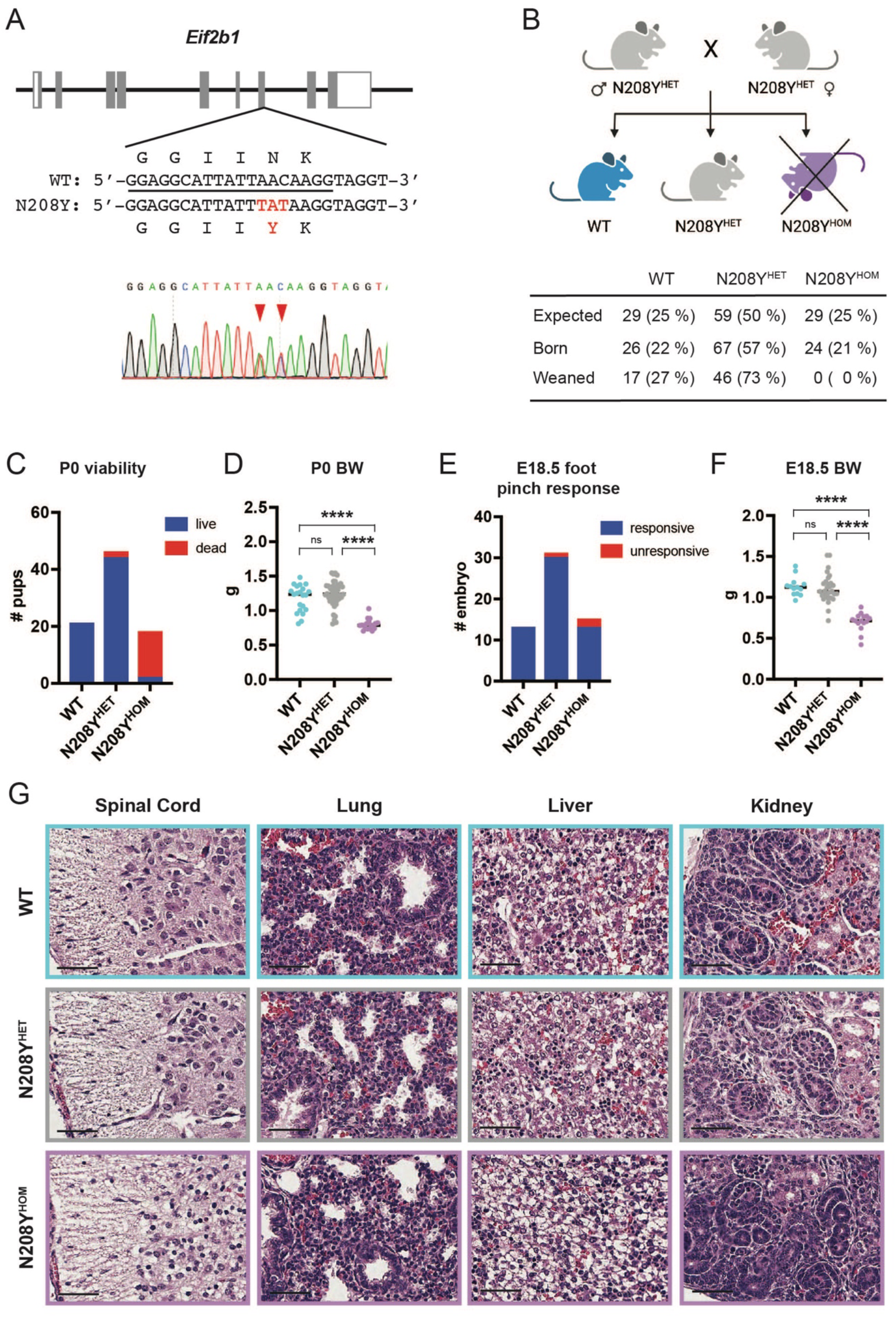
N208Y^HOM^ mice exhibit neonatal lethality with reduced body weight. (A) Schematic representation of mouse *Eif2b1* locus and nucleotide substitution to generate N208Y mutation. Solid gray bars represent open reading frames, Open bars represent UTRs. Underline indicates gRNA sequence used to target the locus. A representative sequencing result of the founder heterozygous mouse is shown. Red arrows indicate nucleotide substitution. (B) Breeding strategy and number of pups born and weaned for each genotype. (C) Number of P0 pups found dead or alive in each genotype. (D) Body weight measurements of WT, N208Y^HET^, and N208Y^HOM^ P0 mice. (C and D) WT, n = 21; N208Y^HET^, n = 46, N208Y^HOM^, n = 18 from 13 litters. (E) Foot pinch responsiveness of E18.5 embryos for each genotype. (F) Body weight measurements of E18.5 embryos for each genotype. (E and F) WT, n = 13; N208Y^HET^, n = 31, N208Y^HOM^, n = 15 from 7 litters. (D and F) Error bars are Standard deviation. One-way ANOVA with Tukey’s multiple comparisons test. **** p < 0.0001, ns = not significant. (G) Representative H & E images of spinal cord, lung, liver, and kidney from E18.5 embryos. scale bars, 50 µm.

To further investigate the early phenotype of N208Y^HOM^ mice, we collected newborn (postnatal day 0, P0) pups and examined their viability and body weight. Whereas the majority of WT and N208Y^HET^ mice were alive, only two out of 18 N208Y^HOM^ newborns were found alive after birth (Figure 2C and Figure Supplement 2A). Body weight measurement revealed that N208Y^HOM^ pups were significantly smaller than the N208Y^HET^ or WT counterparts (Figure 2D). The high number of dead N208Y^HOM^ pups at P0 raised the possibility of stillbirth, rather than neonatal lethality. To distinguish between the two, we isolated embryonic day 18.5 (E18.5) embryos, the terminal embryonic stage right before birth, via Cesarean section of timed pregnant N208Y^HET^ female mice and examined their viability, gross morphology, body weight, and histopathology. We assessed the viability of the embryo via foot pinch test and found that 86.7 % of N208Y^HOM^ embryos (13 out of 15) were alive (Figure 2E and Figure Supplement 2B), suggesting that most N208Y^HOM^ mice are born alive but perish right after birth. Body weight measurement of E18.5 embryos showed N208Y^HOM^ were already significantly smaller than their N208Y^HET^ and WT littermates (Figure 2F). Notably, gross morphological and histopathological examination of N208Y^HOM^ embryos showed normal development of organs (Figure 2G, Figure Supplement 2C, and data not shown).

Taken together, these results indicate that N208Y^HOM^ mice develop normally but have reduced small body weight *in utero* and die quickly after birth due to unknown causes. Thus, this novel mutant mouse model likely recapitulates the most severe forms of VWM which are characterized by early lethality in humans.

### 2BAct rescues the early lethality of N208Y^HOM^ mice significantly extending their lifespan

2BAct is an ISRIB analog with similar potency but with improved solubility and pharmacokinetic properties. We previously demonstrated that this tool compound at a dose of 30 mg/kg in the diet fully attenuates the ISR in the central nervous system (CNS) and prevents neurological defects and loss of myelin in a VWM mouse model caused by the R191H mutation in eIF2Bε (R191H^HOM^) (Wong et al., 2019). Given that ISRIB attenuated the ISR of N208Y eIF2B1 MIN-6 cells (Figure 1G and 1H), we sought to test whether 2BAct could rescue the early lethality of N208Y^HOM^ mice. To this end, we provided control or 2BAct-medicated diets to male and female N208Y^HET^ breeders and maintained their respective diets during the gestational period, nursing period as well as for the offspring after weaning. As expected, the control diet group yielded no N208Y^HOM^ pups upon weaning but, in stark contrast, the group fed with 2BAct had viable N208Y^HOM^ mice which represented 20% of weaned pups (expected mendelian ratio is 25%) (Figure 3A). Given this remarkable rescue, we continued to monitor 2BAct-fed N208Y^HOM^ mice and found that 19 out of 20 were alive at 32 weeks of age and have a 65 % probability of survival at 48 weeks of age (Figure 3B), demonstrating the continued ability of this drug to suppress the deleterious effect of this allele. Taken together, these results demonstrate that 2BAct can rescue the early lethality of N208Y^HOM^ mice and significantly extend their lifespan.

**Figure 3.**
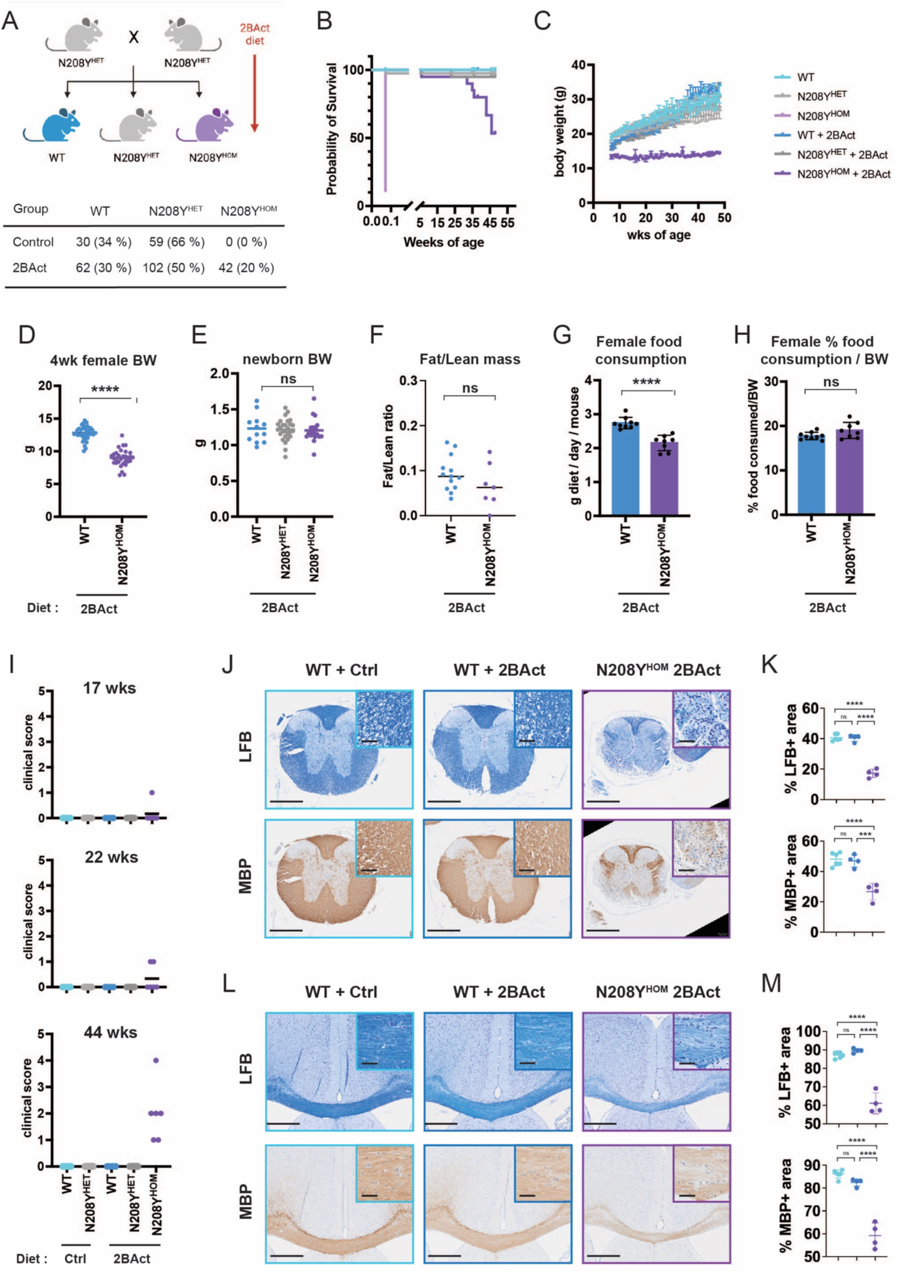
2BAct rescues early lethality of N208Y^HOM^ mice but mice develop VWM pathophysiology with age. (A) Breeding strategy with 2BAct and the number of pups for each genotype after weaning. (B) Kaplan-Meier survival curve. WT, n = 37; N208Y^HET^, n = 80; N208Y^HOM^, n = 18; WT + 2BAct, n = 19; N208Y^HET^ + 2BAct, n = 20, N208Y^HOM^ + 2BAct, n = 20. (C) Body weight measurements of WT, N208Y^HET^, and N208Y^HOM^ mice ± 2BAct. Female WT, n = 3; N208Y^HET^, n = 13; WT + 2BAct, n = 2; N208Y^HET^ + 2BAct, n = 5; N208Y^HOM^ + 2BAct, n = 2. (D) Body weight measurements of 4-week-old female WT and N208Y^HOM^ mice with 2BAct. WT, n = 38; WT + 2BAct, n = 31. (E) Body weight measurement of newborn pups. WT + 2BAct, n = 12; N208Y^HET^ + 2BAct, n = 27, N208Y^HOM^ + 2BAct, n = 20. (A) to (E) Error bars are Standard deviation. Student’s t-test. **** p < 0.0001, ns = not significant. (F) Fat/Lean mass ratio of 3-month-old female mice by EchoMRI. WT + control, n = 8; WT + 2BAct, n = 13; N208Y^HOM^ + 2BAct, n = 7. Female daily food consumption (G) and the percentage of daily food consumption per body weight (H) measured between 5 and 6 weeks of their age. WT + 2BAct, n = 38; N208Y^HOM^ + 2BAct, n = 31. (I) Motor deficit clinical score of WT, N208Y^HET^, and N208Y^HOM^ male and female mice ± 2BAct at 17, 22, and 44 weeks of age. WT, n = 9; N208Y^HET^, n = 27, WT + 2BAct, n = 4, N208Y^HET^ + 2BAct, n = 14; N208Y^HOM^ + 2BAct, n = 6. Representative Luxol Fast Blue (LFB) staining and Myelin Basic Protein (MBP) IHC images of the thoracic region of the spinal cord (J, quantification in K) and the corpus callosum region of the brain (L, quantification in M) from WT+Ctrl, n = 6 (3 females and 3 males); WT+2BAct, n = 4 (2 females and 2 males); N208Y^HOM^+2BAct, n = 4 (2 females and 2 males). Scale bars, 500 µm. Inset scale bars, 50 µm. (K) and (M) Error bars are Standard deviation. One-way ANOVA with Holm-Sidak’s multiple comparisons test. * p < 0.05, ** p < 0.01, *** p < 0.001, **** p < 0.0001, ns = not significant.

### 2BAct-treated N208Y^HOM^ mice show reduced body weight and progressive motor deficits with age

We previously reported that R191H^HOM^ mice have reduced body weight and develop motor deficits with age, and that both phenotypes are fully rescued by preventative treatment with 2BAct (Wong et al., 2019). To evaluate if 2BAct-treated N208Y^HOM^ animals develop any phenotypes with age, we monitored their body weight and motor performance throughout their lifespan. Notably, 2BAct-treated N208Y^HOM^ mice had significantly lower body weight even at the first measured time point at 7 weeks and failed to gain additional weight with age as compared to their N208Y^HET^ and WT littermates (females in Figure 3C and males in Figure Supplement 3A).

The small body size of 2BAct-treated N208Y^HOM^ mice could be due to defects during embryo development and/or the postnatal period associated with increased energy expenditure, reduced fat, and/or decreased food intake. To distinguish amongst these possibilities, we first measured body weight of newborn and 4 weeks old pups treated with 2BAct. While we observed significant body weight reduction of 2BAct-treated N208Y^HOM^ mice at 4 weeks (Figure 3D and Figure Supplement 3B), no differences were detected in newborns across genotypes (Figure 3E). This indicates that 2BAct-treated N208Y^HOM^ mice are born at a normal size and the drug rescued their small body weight during development but not during nursing and the post-weaning period. Next, we measured both fat and lean mass of 3-month-old mice using Echo-MRI. We observed a significant reduction in both fat and lean mass in drug-treated N208Y^HOM^ mice (Figure Supplement 3C), yet no difference in the fat to lean ratio or in the fat to body weight ratio were detected (Figure 3F and data not shown), suggesting that there are no differences in fat mobilization. Finally, we measured daily food consumption and found that 2BAct-treated N208Y^HOM^ mice eat significantly less than their WT counterparts (Figure 3G and Figure Supplement 3D). This difference, however, was lost when food intake was normalized to animal body weight (Figure 3H and Figure Supplement 3E), indicating that increased energy expenditure is not the main cause of reduced weight. These results suggest that 2BAct-treated N208Y^HOM^ mice are born with a normal body weight but then fail to gain weight likely due to reduced food consumption.

In addition to longitudinal tracking of body weight, we also monitored motor performance such as gait and tremor of 2BAct-treated N208Y^HOM^ mice and scored their behavior utilizing the clinical scoring system reported by Traka (2019). We observed that two out of six 2BAct-treated N208Y^HOM^ mice showed mild ataxic phenotypes between 17 and 22 weeks of age (clinical score 1), and by 44 weeks all mutant mice exhibited ataxia with clinical scores ranging from 1 to 4 (2 had mild (score 1), 3 had moderate (score 2), and 1 had severe ataxia (score 4), Figure 3I).

Given that 2BAct-treated N208Y^HOM^ show similar behavioral deficits as untreated R191H^HOM^ VWM mice (Wong et al., 2019), we collected spinal cords and brains from 11-month-old 2BAct-treated N208Y^HOM^ mice and performed histopathological and immunohistochemical analyses. Not surprisingly, spinal cords and brains of N208Y^HOM^ mice showed a significant reduction of the white matter (marked by Luxol Fast Blue (LFB) and myelin basic protein (MBP) staining) in both the spinal cord (Figure 3J and K) and corpus callosum (Figure 3L and M) at this late stage in the disease (only 53.3 % of 2BAct-treated N208Y^HOM^ mice survive to 11 months, Figure 3B). There was also increased inflammation and gliosis (marked by OLIG2, IBA1, GFAP, and ATF3, Figure Supplement 3F-3I) in both tissues, as was shown for R191H^HOM^ mice (Wong et al., 2019).

Collectively, these results suggest that, even though 2BAct is able to rescue the early lethality and significantly extend survival of N208Y^HOM^ mice, drug treatment failed to normalize body weight and did not prevent the development of motor defects and VWM pathology. The phenotype of 2BAct-treated N208Y^HOM^ mice recapitulates that of other VWM mouse models and suggests incomplete rescue of this severe allele.

### 2BAct treatment fully suppresses the ISR in peripheral tissues but not in the central nervous system of N208Y^HOM^ mice

The remarkable rescue of neonatal lethality by 2BAct suggested efficient suppression of the ISR in N208Y^HOM^ mice but the development of neuropathology hinted at chronic ISR activation in the CNS and potentially other tissues. To assess the status of ISR activation, we collected cerebellum, spinal cord and several peripheral tissues from 4-month-old, 2BAct maintained WT or N208Y^HOM^ mice and interrogated the expression of ISR target genes using the nCounter platform that included 95 ISR target genes identified by CLIC (Clustering by Inferred Co-Expression) analysis (Wong et al., 2019) with the addition of *Fgf21*, a stress hormone induced by ISR activation in multiple tissues (Salminen et al., 2017). ISR pathway activation was measured as the change in the average ISR gene panel z-score as compared to 2BAct-treated WT mice, as the treatment with drug did not affect the ISR pathway in WT animals (Figure Supplement 4A). This analysis revealed that the ISR is not significantly induced in any of the peripheral tissues collected (liver, kidney, muscle, spleen, and lung) from 2BAct-treated N208Y^HOM^ mice compared to WT (Figure 4A). In contrast, both the cerebellum and spinal cord showed significant ISR upregulation in 2BAct-treated N208Y^HOM^ mice (Figure 4A). Many known ISR transcriptional targets were induced including ISR-associated transcription factors *Atf5* and *Atf3*, metabolic enzymes and amino acid transporters (Supplemental Table 1).

**Figure 4.**
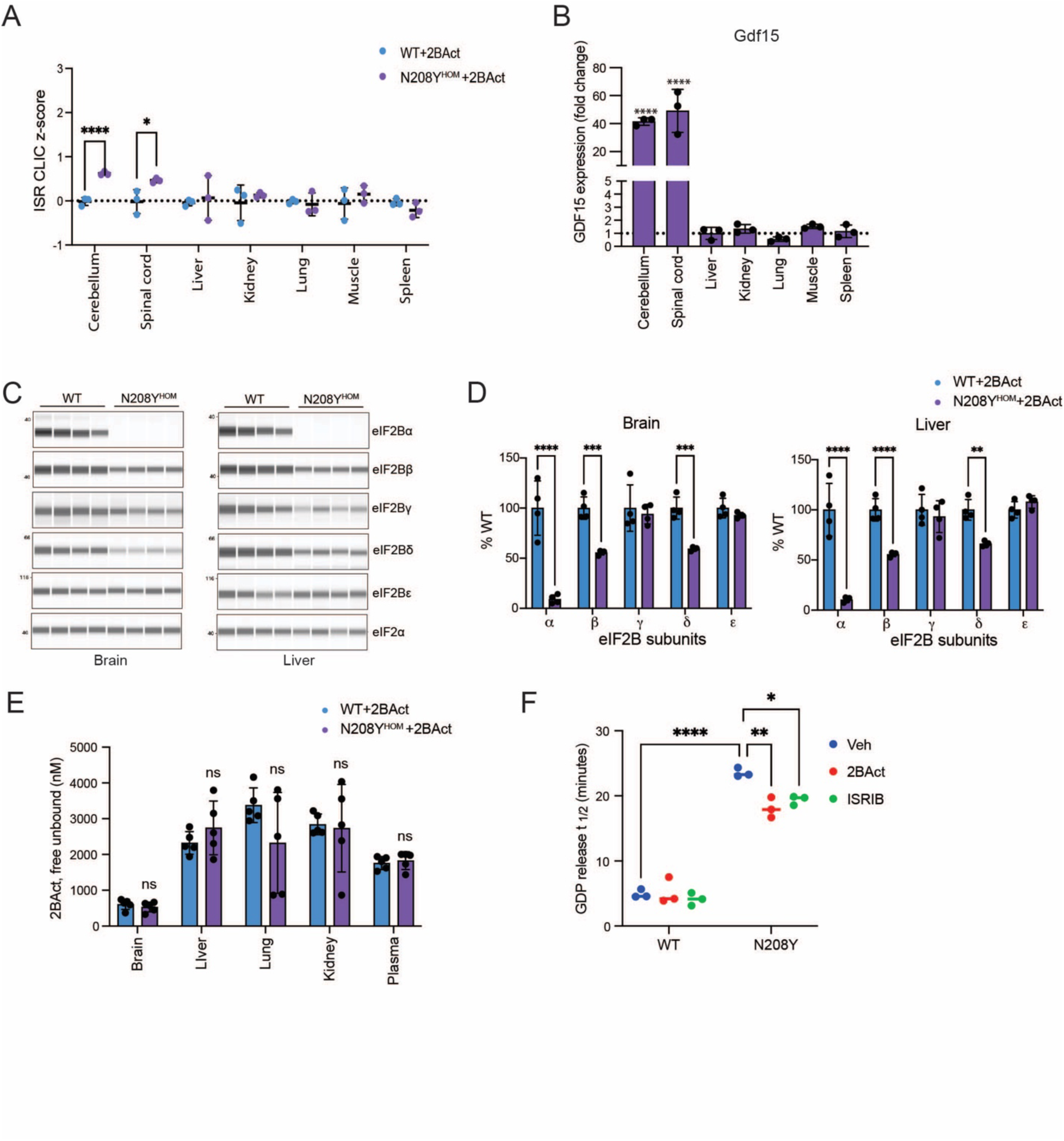
2BAct fully attenuates ISR in peripheral tissues but not in the CNS of N208Y^HOM^ mice. (A) Average z-core of the ISR CLIC genes calculated from nCounter gene expression profiling of various tissues from 4-month-old male mice N208Y^HOM^ + 2BAct normalized to WT + 2BAct. WT + 2BAct, n = 3 and N208Y^HOM^ + 2BAct, n = 3. (B) mRNA expression of *Gdf15* in various tissues normalized to WT + 2BAct using nCounter platform. n=4. Error bars are Standard deviation. Two-way ANOVA with Holm-Sidak’s multiple comparisons test. **** p < 0.0001. (C) Immunoblot of eIF2B subunits in brain (left) and liver (right) lysates from 2BAct maintained 4-month old N208Y^HOM^ male mice. (D) Quantification of bands in (C) normalized to eIF2α expression and represented as % of WT expression. n=4. Error bars are Standard deviation. Two-way ANOVA with Holm-Sidak’s multiple comparisons test. ** p < 0.01, *** p < 0.001, **** p < 0.0001. (E) Tissue drug exposure of 3-month-old male mice. n=5. Error bars are Standard deviation. Two-way ANOVA with Holm-Sidak’s multiple comparisons test. ns, not significant. (F) *In vitro* GEF activity is measured as the half-life of GDP release from eIF2 when added to WT or N208Y MIN-6 cell lysates and is calculated by fitting a single-exponential decay to the Bodipy-FL-GDP fluorescence decay curves. GEF activity was stimulated *in vitro* with 500 nM 2BAct or ISRIB. n=3. Error bars are Standard deviation. Two-way ANOVA with Tukey multiple comparisons test. * p < 0.05, ** p < 0.01, **** p < 0.0001

Notably, expression of *Gdf15*, a known ISR target and secreted protein, was significantly elevated (40-fold increase) in both the cerebellum and spinal cord (Figure 4B). GDF15 is known to impact body weight by reducing food intake through activation of its receptor GFRAL/RET which is expressed only in the brainstem (Wang et al., 2021). The strong upregulation of *Gdf15* transcript in the CNS could lead to reduction in food intake of 2BAct-maintained N208Y^HOM^ mice (Figure 3G and Figure Supplement 3D) and reduced body weight (Figure 3C and Figure Supplement 3A). *Fgf21*, another known ISR target and secreted protein with systemic effects through engagement of its receptor in the brain, was significantly upregulated but only 2-3 fold (Supplemental Table 1).

### The level of eIF2Bα is drastically reduced in all tissues in N208Y^HOM^ mice and results in decreased GEF activity

We were intrigued by the observation that 30 mg/kg 2BAct fully attenuated the ISR CLIC gene signature in the CNS of R191H^HOM^ mice (Wong et al., 2019), but not in N208Y^HOM^ mice. To elucidate the cause of partial suppression of the ISR in the CNS of N208Y^HOM^ mice, we first measured the levels of the eIF2B complex across various tissues. As was the case in ISRIB-maintained N208Y MIN-6 cells (Figure 1B and 1C), Western blot analysis of brain, liver, kidney, and lung lysates showed a drastic reduction in the level of eIF2Bα (> 90% reduction in brain, liver and lung and 70% in kidney). In addition, in mutant tissues, there was a significant reduction in the levels of eIF2Bβ and eIF2Bδ, which together with eIF2Bα form the regulatory sub-complex and the core of the eIF2B decameric complex (Figure 4C, 4D, and Figure Supplement 4B and 4C). The largest reduction in eIF2Bβ and eIF2Bδ was observed in the brain and liver, followed by the lung. In the kidney, these subunits were not significantly reduced.

Given that all eIF2B subunits are essential, it is remarkable that the phenotypes resulting from such a dramatic loss of eIF2Bα across tissues was rescued by 2BAct (neonatal lethality rescue and significant extension of lifespan) as the remaining complex is likely an octamer which is known to have significantly less GEF activity as compared to the full complex (Wong et al., 2018). The similar reduction in eIF2Bα in the brain as compared to liver and lung suggests that this is likely not the main contributor to the partial rescue of the ISR activation in the CNS of N208Y^HOM^ mice.

We then measured drug exposure levels across tissues in both WT and N208Y^HOM^ mice. As previously reported, the level of 2BAct was 3-5 fold lower in the brain than in peripheral tissues or plasma (Wong et al., 2019). Even though we did not observe differences in free unbound 2BAct in CNS between genotypes (Figure 4E), it is possible that the EC_50_ (half maximal effective concentration) of 2BAct is shifted in cells that have reduced eIF2Bα levels and thus, the exposure in the brain may not be sufficient to fully attenuate the ISR in mutant mice. To assess whether cells harboring this mutation have a right-shifted EC_50_, we monitored ATF4 induction in N208Y mutant MIN-6 cells upon removal of ISRIB from the media (Figure 1G) followed by supplementation with varying concentrations of 2BAct, and found that the EC_50_ (27.8nM) (Figure Supplement 4D) was comparable to the one previously reported in wild type cells (33 nM)(Wong et al., 2019). Importantly, the level of unbound 2BAct in the brain of mutant mice was 19-fold higher than this EC_50_, and thus, a predicted saturating concentration.

Given that this level of 2BAct did not attenuate the ISR in the CNS, we then measured the maximal GEF activity of the endogenous eIF2B complex attained at 2BAct-saturating conditions in lysates from N208Y or WT MIN6 cells. To this end, we added fluorescently labeled GDP-loaded eIF2 substrate to lysates and determined the rate of reduction in fluorescence which corresponds to the rate of GDP release from eIF2. As shown in Figure 4F, lysates from mutant cells show a dramatic, although not unexpected, reduction in GEF activity (slower release) as compared to WT lysates (23.6 min +/-0.681 vs 4.9 min +/-0.649). Both ISRIB and 2BAct at saturating concentration (500 nM) boosted the GEF activity of N208Y lysates but the rate of GDP release is improved to a small extent as compared to the rate of WT lysates.

Collectively these results indicate that induction of the ISR in the brain and spinal cord is likely due to reduced eIF2B GEF activity in 2BAct-treated N208Y^HOM^ mice and differential sensitivity of the CNS to mutated eIF2B alleles as is observed in other VWM mouse.

### 2BAct withdrawal leads to rapid weight loss and motor deficits

Together, the *in vitro* and *in vivo* data demonstrated that N208Y mutant cells and N208Y^HOM^ mice require the presence of an eIF2B activator (ISRIB or 2BAct) to sufficiently boost GEF activity in order to grow or survive. To assess the physiological consequence of drug removal, we switched a subset of WT and N208Y^HOM^ mice from 2BAct-medicated diet to control diet at 3 to 4 months of age and monitored body weight and motor performance daily (Figure 5A). No differences in body weight were observed amongst the four groups up to 3 days after 2BAct withdrawal. However, 2BAct-withdrawn N208Y^HOM^ mice showed rapid decline in their body weight starting at day 4, losing more than 15% of their body weight within the next two days (males in Figure 5B and females in Figure Supplement 5A). We also examined their gait and scored their behavior throughout the washout period. Similar to the body weight measurements, mice in all four groups did not exhibit any obvious abnormalities up to day 3.

**Figure 5.**
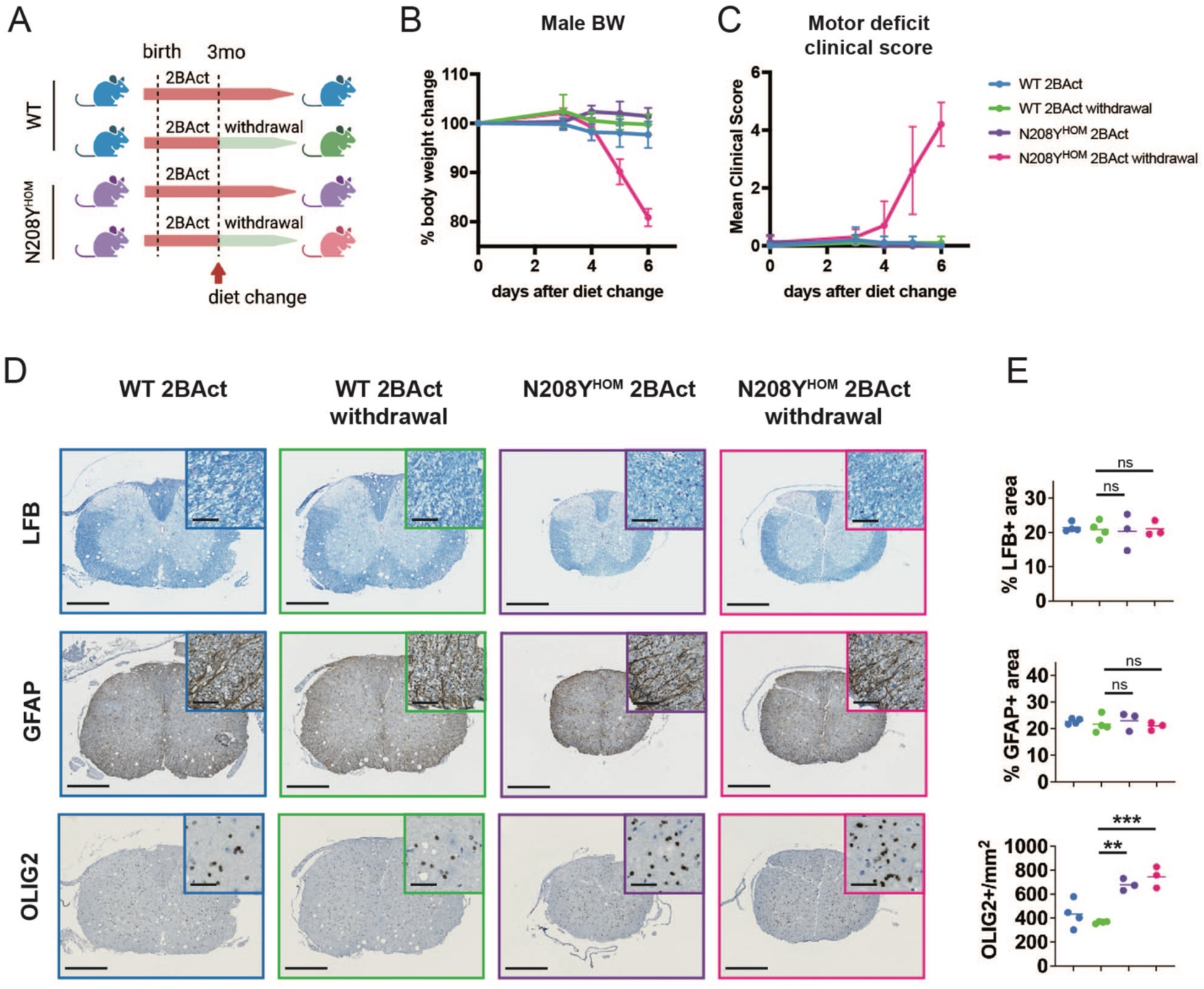
Withdrawal of 2BAct leads to rapid weight loss and motor deficit. (A) Schematic representation of 2BAct withdrawal strategy of 2BAct-rescued WT and N208Y^HOM^ mice. Percentage of body weight change (B) and mean motor deficit clinical score (C) of 4-month-old male mice after 2BAct withdrawal. For clinical scoring, gait, tremor, and other motor deficits were assessed according to the published classification (Traka, 2019). WT + 2BAct, n = 5; WT + 2BAct withdrawal, n = 5; N208Y^HOM^ + 2BAct, n = 4; N208Y^HOM^ + 2BAct withdrawal, n = 5. (D) Luxol fast blue and Immunohistochemical staining using antibodies specific to GFAP, and OLIG2 from spinal cords of 3-month-old females after 7-day 2BAct withdrawal. WT + 2BAct, n = 4; WT + 2BAct withdrawal, n = 4; N208Y^HOM^ + 2BAct, n = 3; N208Y^HOM^ + 2BAct withdrawal, n = 3. Scale bar 500 µm and 50 µm (inset) (E) Quantification results of (D). One-way ANOVA with Holm-Sidak’s multiple comparisons test. ** p < 0.01, *** p < 0.001, ns = not significant.

After day 4, 2BAct-withdrawn N208Y^HOM^ mice developed ataxia and tremor, and these phenotypes worsened over time (Figure 5C). Given the rapid onset of motor deficit upon withdrawal, we wondered whether accelerated demyelination could partially explain the rapid deterioration seen upon drug removal. To measure myelin content, we performed histological and immunohistochemical analyses of the spinal cords from 3-month-old female mice after 7 days of 2BAct withdrawal. Similar to the 11 month old 2BAct-maintained N208Y^HOM^ cohort (Figure 3J and 3K and Figure Supplement 3F and 3G), there was an increase in OLIG2-positive oligodendrocyte lineage cell number at 4 months in the 2BAct-maintained group (Figure 5D and 5E) which likely represents early stages of disease progression. However, no differences in myelination (LFB staining) were observed at this earlier time point. After drug withdrawal, we did not detect any change in GFAP, OLIG2 or myelin staining, suggesting that the onset of motor deficit is likely not due to accelerated demyelination.

The striking onset of body weight loss and motor deficits upon drug removal clearly demonstrate that 2BAct is required to maintain a physiological state that is compatible with life in N208Y^HOM^ mice.

### 2BAct withdrawal triggers ISR induction in all tissues tested in N208Y^HOM^ mice

The rapid effects of drug withdrawal in N208Y^HOM^ mice suggested that 2BAct efficiently suppresses the ISR. To broadly survey ISR status, we collected both peripheral and CNS tissues as well as plasma at 0 or 3 days after switching to a control diet (Figure 6A), and measured ISR CLIC gene expression using the nCounter platform. The average ISR CLIC z-score showed further activation of the stress signature after withdrawal in both the cerebellum and the spinal cord in N208Y^HOM^ mice as compared to mutant animals kept on 2BAct diet (Figure 6B). In contrast, no significant change in ISR gene expression was observed in 2BAct-withdrawn WT mice when compared to their 2BAct-maintained counterparts.

**Figure 6.**
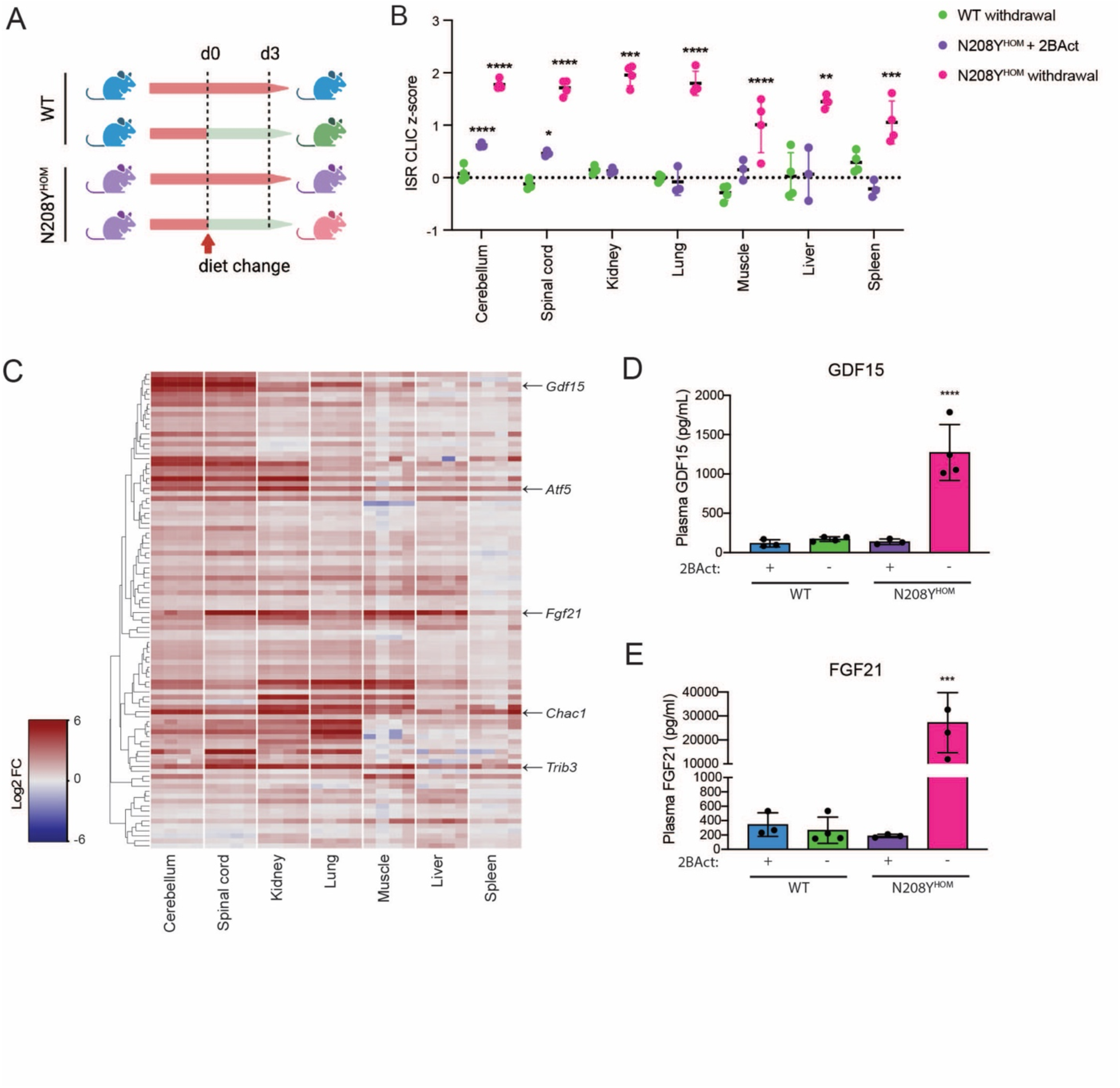
2BAct withdrawal rapidly induces plasma and tissue ISR signatures. (A) Schematic representation of tissue and plasma collection 3 days after 2BAct withdrawal. (B) Average z-core of ISR CLIC genes calculated from nCounter gene expression profiling with 2BAct withdrawal in cerebellum, spinal cord, kidney, liver, lung, muscle and spleen from N208Y^HOM^ 4-month-old male mice normalized to WT 2BAct maintained littermates. Error bars are standard deviation. Two-way ANOVA with Holm-Sidak’s multiple comparisons test. ** p < 0.01, *** p < 0.001, **** p < 0.0001. (C) Heatmap of ISR CLIC genes using nCounter gene expression platform for all tissues from N208Y^HOM^ 4-month old male mice after 2BAct withdrawal normalized to WT 2BAct maintained littermates. ELISA quantification of plasma GDF15 (D) and FGF21 (E) levels 3 days after 2BAct withdrawal from 4-month-old male WT and N208Y^HOM^ mice. One-way ANOVA with Holm-Sidak’s multiple comparisons test. ** p < 0.01, *** p < 0.001 **** p < 0.0001. (B and C) WT + 2BAct, n = 3; WT + 2BAct withdrawal, n = 4; N208Y^HOM^ + 2BAct, n = 3; N208Y^HOM^ 2BAct withdrawal, n = 4

We next measured the impact of drug withdrawal in peripheral tissues, all of which did not have a measurable ISR in 2BAct-maintained N208Y^HOM^ mice. Notably, we observed a significant increase in the average ISR CLIC z-score upon diet change in all peripheral tissues of N208Y^HOM^ mice (Figure 6B). The fold increase after a 3 day drug washout was comparable across cerebellum, spinal cord, kidney and lung but smaller in muscle, liver and spleen. As shown in the heatmap in Figure 6C, we observed heterogeneity in the pattern of ISR CLIC genes induced across the tissues surveyed. *Atf5*, *Chac1* and *Trib3* are three known ISR targets that were significantly elevated in all tissues in N208Y^HOM^ after withdrawal (Figure 6C and Supplemental table 2). *Gdf15* was highly induced in both CNS tissues (the spinal cord and the cerebellum) but to a lesser extent in the peripheral tissues. In contrast, *Fgf21* was highly upregulated in the spinal cord (> 98 fold) but modestly in the cerebellum (>7 fold) and was also significantly induced in all peripheral tissues upon removal of 2BAct. The unique pattern of ISR target gene induction likely reflects the heterogeneity in the expression of transcriptional co-activators and co-repressors that modulate the impact of ISR-dependent transcription factors such as ATF4 across tissues. We next assessed the impact of 2BAct withdrawal on the accumulation of GDF15 and FGF21 in plasma and observed a significant increase 3 days after drug withdrawal in N208Y^HOM^ mice (Figure 6D and 6E). Moreover, no difference in levels of these circulating proteins was detected between 2BAct-maintained N208Y^HOM^ and WT mice. This is in agreement with the lack of ISR CLIC gene induction in peripheral tissues of 2BAct-maintained mutant mice.

Taken together, these results demonstrated that 2BAct fully suppresses the ISR in peripheral tissues of N208Y^HOM^ mice and that, upon washout, the stress response is activated across all measured tissues leading to fast accumulation of GDF15 and FGF21 in the plasma. This underscores the importance of continuous access to an eIF2B activator to suppress induction of this stress response to sustain the life of this mutant mouse strain.

### The transcriptome of 2BAct-maintained N208Y^HOM^ and untreated R191H^HOM^ mice showed a high level of similarity and drug removal elicited extensive gene expression changes

2BAct-treated N208Y^HOM^ mice exhibited progressive motor deficits and demyelination with age that phenocopied disease progression of untreated R191H^HOM^ mice (Figure 3I-3M)(Wong et al., 2019). Moreover, upon 2BAct withdrawal, N208Y^HOM^ mice showed rapid onset of severe motor deficits and pervasive activation of the ISR. To compare the gene expression changes in the CNS of these two VWM mouse models in an unbiased fashion, we performed bulk RNA-seq analysis of cerebellum from 4 month-old N208Y^HOM^ mice before and after drug removal and from untreated 5-month old R191H^HOM^ mice. Differential expression analysis comparing each VWM mutant to its WT control showed a greater number of significantly differentially expressed genes in the cerebellum of 2BAct-treated N208Y^HOM^ (n = 710 with log_2_ fold change > 0.5 and q < 0.05) than in untreated R191H^HOM^ (n =210) (Figure 7A and Supplemental Table 3). 2BAct did not affect gene expression in WT mice underscoring its high specificity (Figure Supplement 7A). ISR target genes were upregulated in both mutant mice, but the effect sizes were greater in 2BAct-treated N208Y^HOM^ mice (red circles in Figure 7A and 7B). Overall, fold changes for genes significant in either the R191H^HOM^ vs. WT comparison or the 2BAct-treated N208Y^HOM^ vs. WT comparison were well correlated (p=7.40 x 10^-240^, R^2^=0.65, Figure 7B), indicating molecular similarity between the two VWM mutant mouse models.

**Figure 7.**
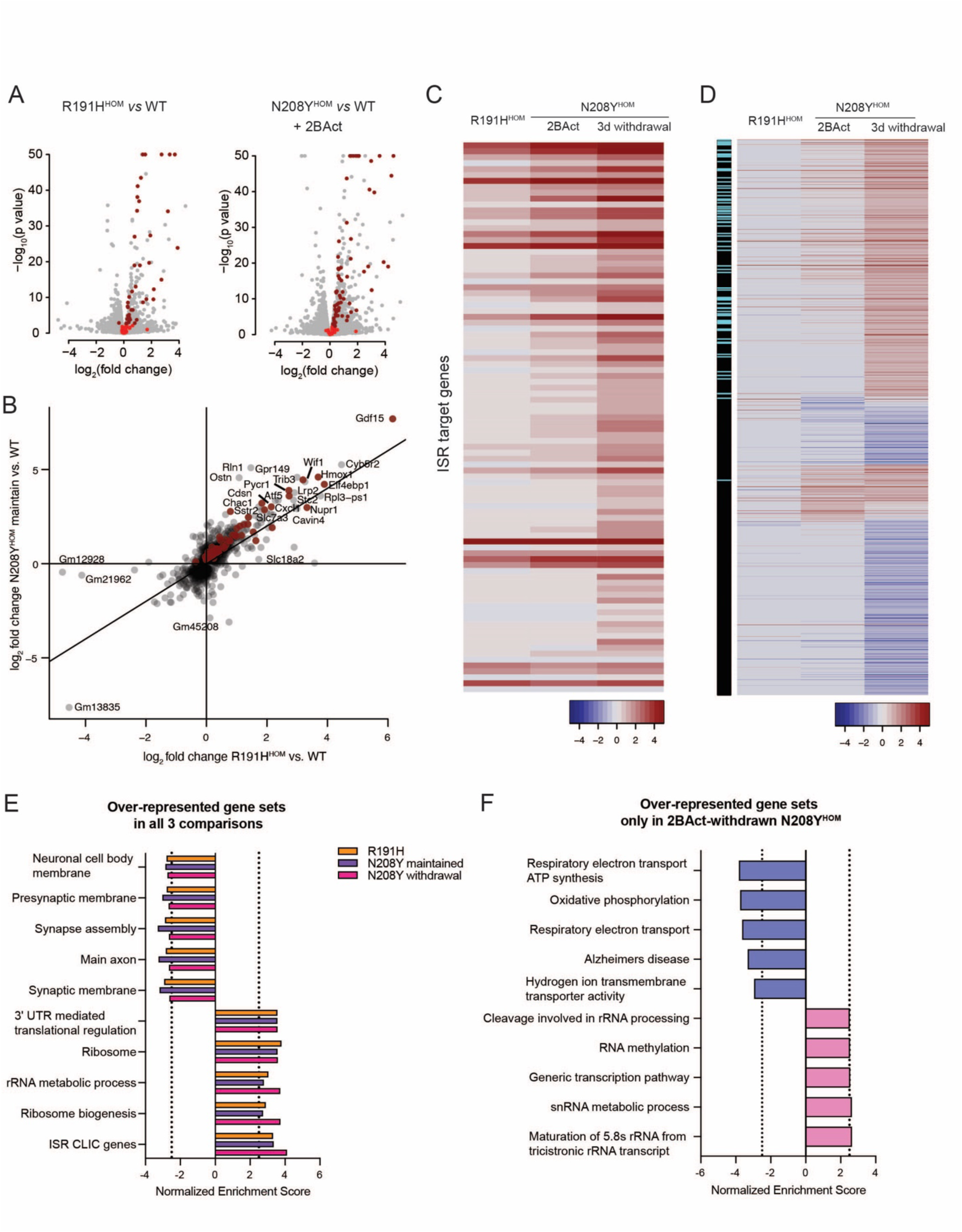
The cerebellum transcriptome of 2BAct-maintained N208Y^HOM^ mice showed similarity to that of R191H^HOM^ mice and drug removal elicited extensive changes in gene expression. (A) Volcano plots displaying gene expression differences in gene expression between R191H^HOM^ and WT (left) and N208Y^HOM^ + 2BAct and WT + 2BAct (right). Red dots show ISR CLIC genes and dark red indicate significantly upregulated ISR CLIC genes (adj p < 0.05). (B) Correlation plot showing differentially expressed genes from either R191H^HOM^ or N208Y^HOM^ + 2BAct compared to their respective wild type groups. The genes with more than 3 log_2_ fold change were labeled. Dark red dots indicate significantly upregulated ISR CLIC genes (adj p < 0.05). Heat map of (C) ISR CLIC genes and (D) all significantly differentially expressed genes from R191H^HOM^ vs WT, N208Y^HOM^ + 2BAct vs WT + 2BAct, and N208Y^HOM^ 2BAct withdrawal vs WT + 2BAct comparisons. In (D), on the left, ISR CLIC genes are shown in light blue. The top 5 most significantly overrepresented gene sets in all three comparisons (E) or only in N208Y^HOM^ 2BAct withdrawal vs WT + 2BAct comparison (F). Cutoff of Normalized Enrichment Score (2.5) is presented as dotted lines.

In agreement with our prior analysis (Figure 6C), withdrawal of 2BAct led to a further increase in ISR CLIC gene expression in the cerebellum of N208Y^HOM^ mice (adj p = 0.0013, Figure 7C). In addition, we observed significant differential expression of many additional genes (1733 genes with log_2_ fold change > 0.5 and q < 0.05) and gene sets in 2BAct-withdrawn N208Y^HOM^ mice (Figure 7D and Supplemental Table 4). Among these, we observed gene sets that, like ISR CLIC genes, were upregulated in all three comparisons to WT (R191H^HOM^, 2BAct-maintained N208Y^HOM^, and 2BAct-withdrawn N208Y^HOM^). These gene sets were related to ribosome biogenesis, 3’UTR mediated translational regulation and rRNA metabolic process (Figure 7E and Supplemental Table 4). Similarly, gene sets related to neuronal synapse structure or function were downregulated in all three comparisons. Unique gene sets that changed only in 2BAct-withdrawn N208Y^HOM^ included downregulation of gene sets related to oxidative phosphorylation, respiratory electron transport, and gluconeogenesis (Figure 7F, Supplemental Table 4).

In summary, the transcriptomic analysis of the cerebellum further supports the similarity between 2BAct-maintained N208Y^HOM^ and untreated R191H^HOM^ mice. Partial suppression of the ISR by 2BAct in the CNS of N208Y^HOM^ mice results in gene expression changes that are similar to those observed in the milder R191H VWM model. Furthermore, this analysis revealed that drug withdrawal leads to extensive remodeling of the transcriptome in the cerebellum, likely reflecting activation of the ISR across additional cell types and ensuing downstream transcriptional effects. Broad transcriptional changes in the CNS, together with rapid activation of the ISR in peripheral tissues, likely drive the rapid physiological decline observed in N208Y^HOM^ mice when drug is removed.

## Discussion

eIF2B is a decameric complex that is essential for initiation of mRNA translation in all eukaryotic cells. Its enzymatic activity is required for GTP exchange of eIF2 and delivery of the initiator methionine tRNA to the ribosome to initiate protein synthesis. Importantly, eIF2B is a stress sensor and key ISR effector that is inhibited by phosphorylation of its substrate eIF2 eliciting a brake in protein synthesis and activation of the ISR transcriptional response. We recently found that eIF2B complex stability and activity is also directly modulated by sugar phosphates (Hao et al., 2021). We hypothesized that this is a potential mechanism for direct coupling of protein synthesis rate, a costly cellular process, to the energy status in the cell. As suggested by its homology to an archeal sugar metabolizing enzyme, F6P and related metabolites bind to the ancestral site in eIF2Bα and this binding promotes formation of the decameric eIF2B complex, which is the most active enzyme.

To determine the cellular consequences of eIF2B metabolite binding, we generated MIN-6 cells carrying either of two point mutations, E198K and N208Y, that decrease or abolish sugar phosphate binding, respectively, and found a striking reduction in eIF2Bα subunit levels and GEF lysate activity. Moreover, their generation required supplementation with the eIF2B activator ISRIB, underscoring the ability of this compound to boost GEF activity in conditions of severely limited eIF2Bα dimer. The observed higher intrinsic instability of purified E198K and N208Y eIF2Bα and lack of stabilization upon F6P addition suggests that sugar phosphates act as obligate cofactors for eIF2Bα stability in cells. We hypothesize that limiting the amount of eIF2Bα, mediated by sugar phosphate binding pocket occupancy, acts as a failsafe mechanism that sustains the brake in protein synthesis under prolonged nutrient deprivation independent of the phosphorylation status of eIF2. Notably, N209Y eIF2Bα (N208Y in humans and mice) yeast cells also show dramatically reduced eIF2Bα levels (Richardson et al., 2004), demonstrating that the impact of sugar phosphate binding on subunit stability is highly conserved.

Given the dramatic cellular effect of the N208Y eIF2Bα mutation in mammalian cells, we were intrigued by the association of this sugar phosphate binding mutant with VWM, a recessive and rare leukodystrophy caused by partial loss of function alleles in any of the five subunits of eIF2B. To date, more than 300 mutations have been associated with VWM (Hamilton et al., 2018) and as is often the case, the N208Y eIF2Bα carrying patient was a compound heterozygote. The second allele is a substitution predicted to impair a splicing donor nucleotide leading to a frame shift and production of a severely truncated eIF2Bα protein (van der Knaap et al., 2002). This patient was diagnosed with VWM at the age three and reported to survive for 11 years (Slynko et al., 2021). Not surprisingly, and in contrast to previously reported VWM mouse models that show progressive loss of white matter with age, N208Y^HOM^ mice showed neonatal lethality. This is the earliest and most severe phenotype of a VWM-associated mutation reported to date and more akin to severe human VWM cases that present with antenatal and neonatal lethality. Surprisingly, besides their reduced body weight, we found no obvious defects in E18.5 embryos and newborn pups. Gross and microscopic examination showed that all organs developed normally with no obvious pathological features. This was a surprising finding given the proposed role of ISR activation in cell fate determination in various tissues in mice (Heijmans et al., 2013; Scheuner et al., 2001; van Galen et al., 2018; Zhang et al., 2006; Zismanov et al., 2016). Why do N208Y^HOM^ mice die right after birth? A deeper investigation is required to identify the cause of death. The observed phenotype in a small number of animals, responsiveness to foot pinching but no sign of breathing right after birth, suggest that breathing failure could be a likely cause. Breathing upon birth requires proper lung function as well as neuromuscular coordination controlled by neurons in the brain and spinal cord that regulate respiratory muscles (Turgeon and Meloche, 2009). Therefore, certain cell types that comprise the neuromuscular lung system may be impacted during development, but their effects may not be evident in pathological examinations of mutant mice.

Our prior success with preventive 2BAct treatment in R191H^HOM^ mice, a well characterized VWM mouse model that carries an early onset human mutation in the ε subunit, revealed the beneficial impact of this mechanism of action in this devastating disease. The ability of 2BAct to rescue the neonatal lethality of N208Y^HOM^ mice underscores the remarkable efficacy of these molecules in a disease driven by eIF2B hypomorphic alleles. N208Y^HOM^ mice are particularly interesting as this model is characterized by a dramatic decrease in eIF2Bα, an essential subunit that promotes complex formation. Importantly, 2BAct sufficiently boosted the GEF activity of the remaining eIF2B complex *in vivo* to significantly extend the lifespan of mutant mice beyond 11 months.

Although early lethality was rescued by 2BAct, drug-maintained N208Y^HOM^ mice showed reduced body weight and developed VWM pathology with age, phenocopying untreated R191H^HOM^ mice, which live up to 10 months (Dooves et al., 2016). We found that the ISR was not fully suppressed and restricted to the CNS in 2BAct-maintained N208Y^HOM^ mice and unbiased transcriptome analysis of the cerebellum confirmed its similarity to the transcriptome of untreated R191H^HOM^ mice. Importantly, 2BAct fully suppressed the ISR in peripheral tissues. Why do 2BAct-rescued mice have an ISR only in the CNS? eIF2Bα levels were consistently and significantly reduced across tissues suggesting that differential stability is unlikely to explain the difference in response across tissues. We previously showed that ISRIB can boost the GEF activity of purified eIF2Bβγδε but it does not reach the level of activity of the untreated full decameric eIF2B complex (Wong et al., 2018). This is consistent with the modest increase in GEF activity in MIN6 mutant lysates upon 2BAct or ISRIB addition. In K562 cells, however, ISRIB can attenuate ISR induction triggered by rapid and complete eIF2Bα degradation (Schoof et al., 2021). Moreover, ISRIB suppressed ATF4 induction and ISR target genes in N208Y MIN6 cells. These cell-based results, in conjunction with the observed lack of ISR activation in peripheral tissues of 2BAct-treated N208Y^HOM^ mice, demonstrate that a small boost of GEF activity by ISRIB-like compounds can fully attenuate ATF4 production and ISR target gene expression. Intriguingly, astrocytes are particularly susceptible to eIF2B hypomorphic mutations showing preferential activation of the ISR in R191H^HOM^ mice (Wong et al., 2019). Preferential ISR activation in glial cells has also been reported in VWM patients (Abbink et al., 2019). Given the CNS restricted susceptibility observed in R191H^HOM^ mice, it is not surprising that partial rescue of the GEF activity of N208Y mutant cells by 2BAct results in a similarly restricted pattern of ISR activation.

2BAct withdrawal triggered significant induction of ISR in all tissues in N208Y^HOM^ mice and led to high levels of circulating FGF21 and GDF15, two potent circulating ISR target proteins with broad systemic effects. Mice quickly deteriorated, demonstrating the need for access to 2BAct once treatment is initiated to support life. Unlike pleiotropic stress-induced animal models, induction of the ISR response in N208Y^HOM^ mice is a consequence of a reduction in eIF2B activity upon drug withdrawal and provides a unique genetic model to temporally control ISR activity *in vivo* in the absence of activation of parallel pathways commonly triggered by other insults. Bulk RNA-seq analysis of N208Y^HOM^ cerebellum revealed that 2BAct withdrawal leads to the rapid downregulation of gene sets related to oxidative phosphorylation, respiratory electron transport and ATP synthesis (Figure 7F). This is not surprising given the further reduction in GEF activity that cells experience upon washout of 2BAct which likely result in a decrease in the rate of protein synthesis and energy demand. Notably, ISR activation led to tissue-specific differences in expression of target genes that comprise the ISR signature. Single cell analysis of various tissues is likely to reveal a complex transcriptional network and allow identification of transcription factors that contribute to ISR target gene expression. Therefore, this new genetic model provides us with a tool to tune pathway activation by controlling the dose and availability of 2BAct and thus allow deeper exploration of temporal activation of the ISR pathway across cell types and tissues.

## Materials and Methods

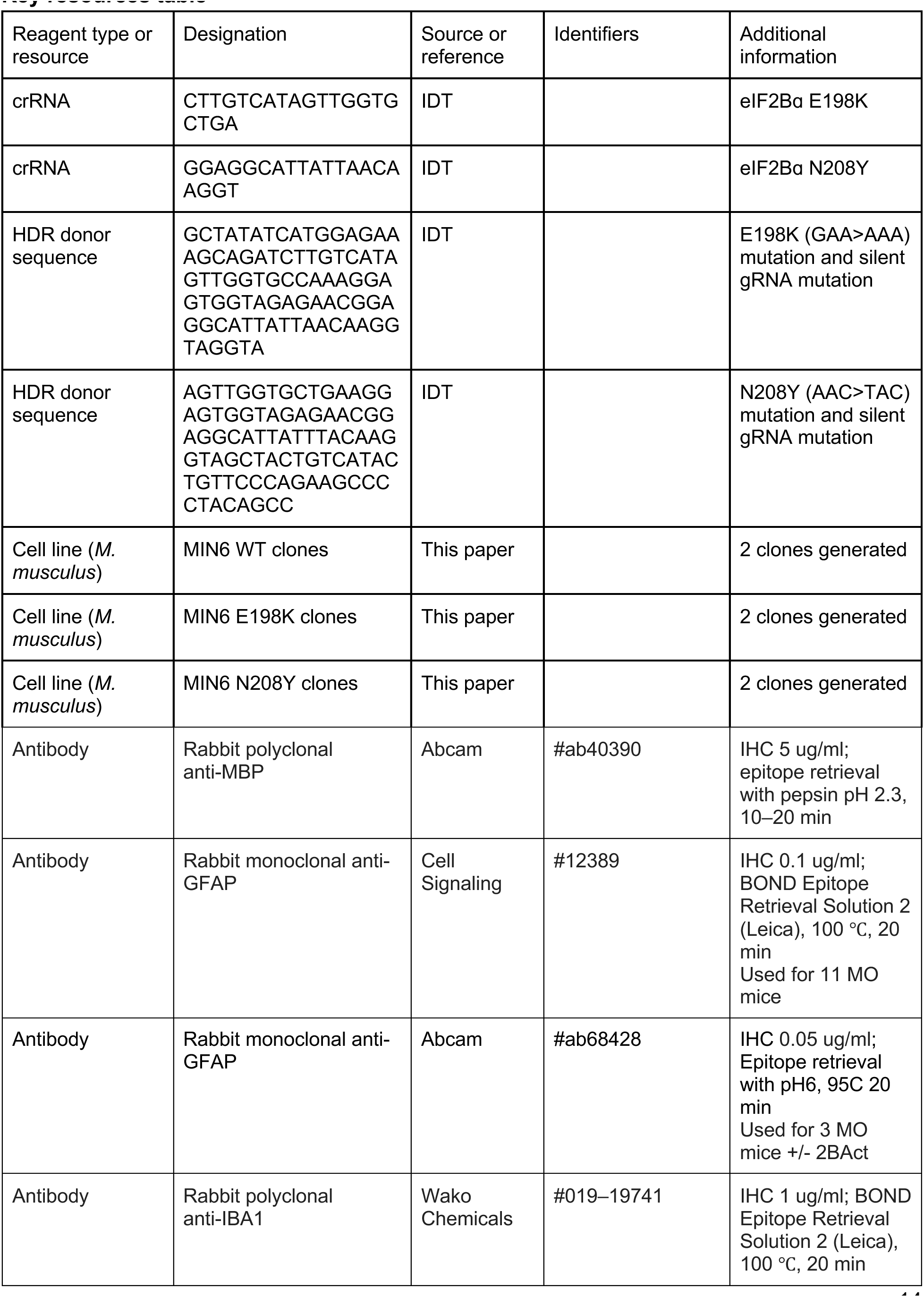
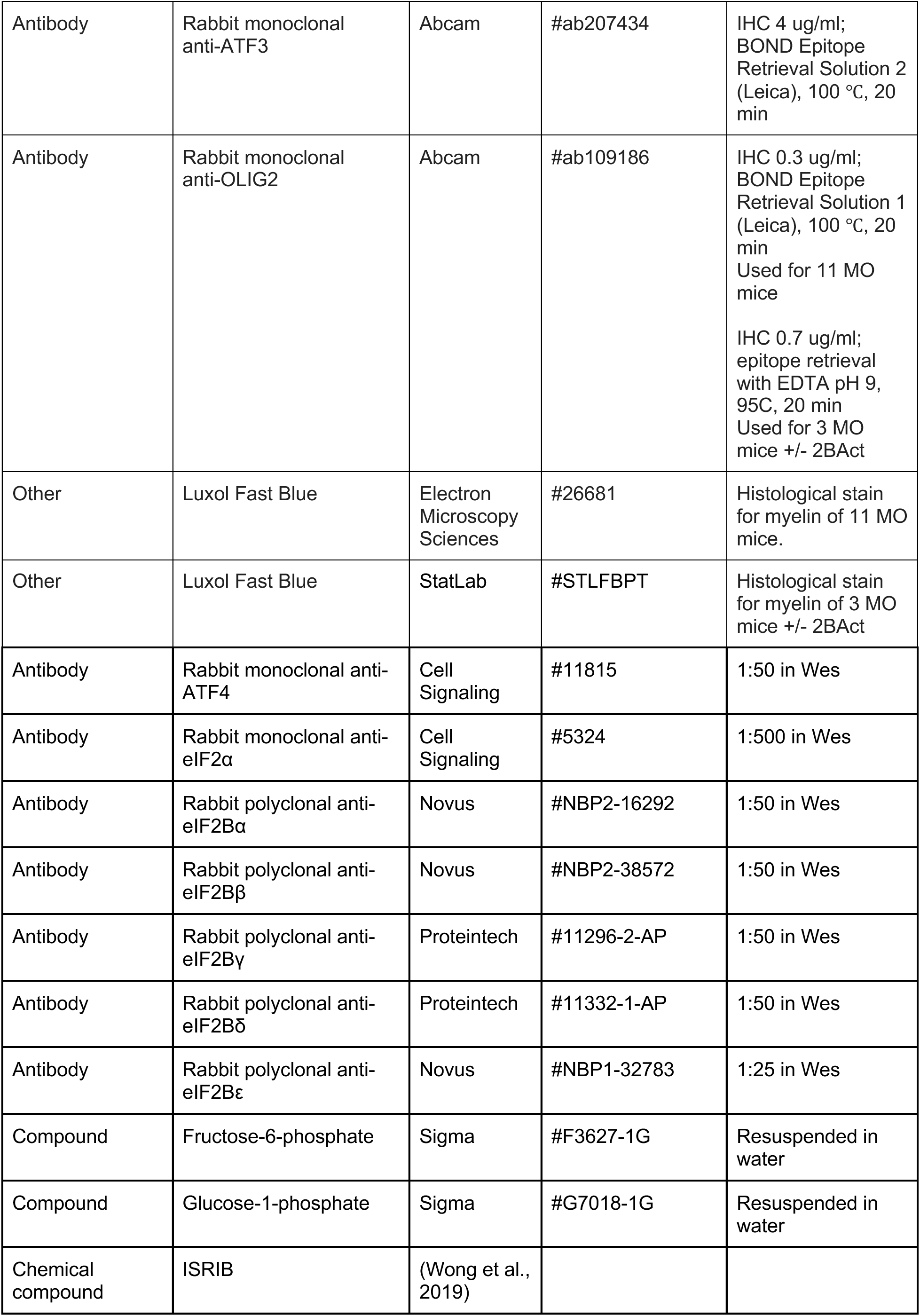
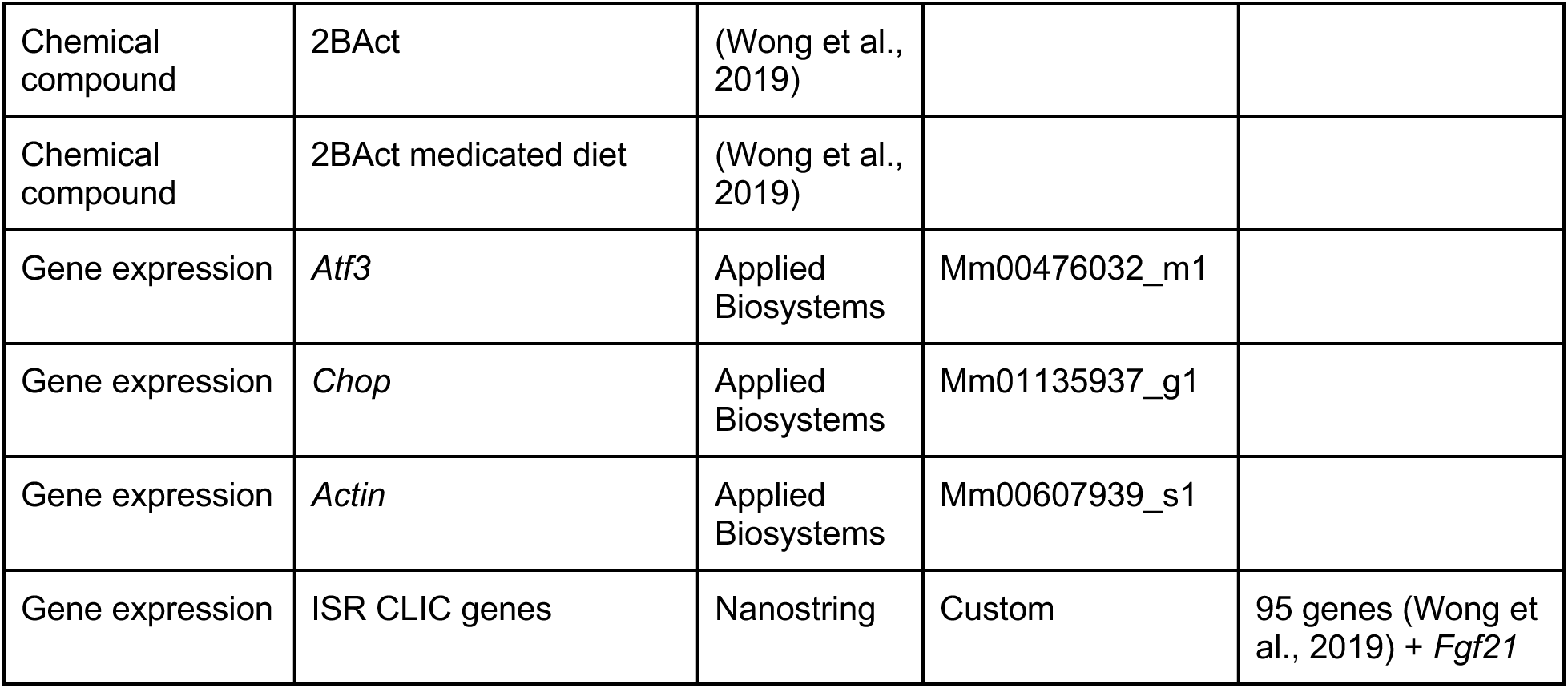
Key resources table

### Generation of mutant cell lines and cell culture

The eIF2Bɑ E198K and N208Y mutations were introduced into MIN-6 cells (AddexBio #C0018008) using the Alt-R CRISPR-Cas9 System (Integrated DNA Technologies, Coralville, IA, USA) following manufacturer’s protocol. The RNP and HDR donor was delivered to MIN-6 cells by electroporation using the Lonza 4D-Nucleofector (Lonza Group AG, Basel, Switzerland), using the 96-well SE reagent kit and program CM-150. Nucleofected cells were incubated in media containing Alt-R HDR Enhancer and 200 nM ISRIB for 24 hr before media was changed to complete media + ISRIB. After 48 hr, cells were single-sorted into 96-well plates containing complete media + ISRIB using a BD Biosciences FACSAria Fusion (BD Biosciences, San Jose, CA, USA). Clones were expanded and genomic DNA was sequenced to confirm the desired mutations. Experiments involving ISRIB withdrawal from cell lines were accomplished by one round of PBS wash followed by addition of complete media without ISRIB.

MIN-6 cell lines (AddexBio #C0018008) were grown in complete media (DMEM containing 25 mM glucose supplemented with 15% FBS, 100 U/mL streptomycin, 100 U/mL penicillin sulfate and 75 uM β-mercaptoethanol). Cell lines were maintained in complete media + 200 nM ISRIB. Cell lines were incubated at 37°C, 5% CO2.

### Immunoblots

Cell lysates were prepared in RIPA buffer (Thermo Scientific #89900) with protease/phosphatase inhibitors (Thermo Scientific #78444). Cells were lysed on ice for 10 minutes then centrifuged (21,000 x g, 15 min, 4C) to remove cellular debris. Tissue lysates were homogenized by mortar pestle grinding in liquid nitrogen prior to bead-based homogenization in RIPA buffer. All lysates were incubated on ice for 10 minutes then centrifuge (21,000 x g, 15 min, 4C) to remove cellular debris. Lysate concentration was determined using BCA and cell protein lysate samples were adjusted to 0.5 mg/mL and tissue protein lysate samples were adjusted to 0.75 mg/mL. Samples were run on a ProteinSimple Wes system (Bio-Techne, Minneapolis, MN, USA) using a 12-230 kDa separation module. GraphPad Prism (La Jolla, CA, USA) was used to perform statistical analyses utilizing two way ANOVA with Holm-sidak performed post-hoc to correct for multiple comparisons.

### Differential scanning fluorimetry

Recombinant eIF2Bα WT, E198K, N208Y were purified as previously described (Hao et al., 2021). Thermal shift assay was performed on Prometheus Panta NT.48 from NanoTemper technologies by measuring the intrinsic dual-UV fluorescence change in tryptophan and tyrosine residues in proteins at emission wavelengths of k = 330 and 350 nm.The ratio of the recorded emission intensities (Em350nm/Em330nm), which represents the change in TRP fluorescence intensity as well as the shift of the emission maximum to higher wavelengths (“red-shift”) or lower wavelengths (“blue-shift”) was plotted as a function of the temperature. The fluorescence intensity ratio and its first derivative were calculated and determined to be the melting temperature (Tm), with the manufacturer’s software (PR.Panta Control and PR.Panta Analysis). The samples were loaded using capillaries in a volume of 10µL on Prometheus Panta NT.48 from NanoTemper Technologies. 50 µM eIF2Bα was incubated with 2-fold serial dilution of the indicated sugar phosphates at a concentration titration of 0.046mM - 3mM, and then subjected to thermal change from 25°C - 85°C with a ramp rate of 1°C/min.

### Generation of *Eif2b1* (p.N208Y) mutant mouse model

The mutant mouse strain carrying the *Eif2b1* p.N208Y point mutation was generated as a service by Cyagen Inc. through targeted mutagenesis in mouse embryos using the CRISPR/Cas9 system. In brief, the Cas9 protein, the synthesized sgRNA (5’-GGAGGCATTATTAACAAGGTAGG-3’) and ssDNA donor (5’-GAGAAAGCAGATCTTGTCATAGTTGGTGCTGAAGGAGTGGTAGAGAACGGAGGCAT TATTTATAAGGTAGGTACTGTCATACTGTTCCCAGAAGCCCCTACAGCCTGAGCAAG ACCTTGTCAG-3’) harboring the p.N208Y mutation (AAC to TAT) were co-injected into the cytoplasm of pronuclear stage embryos. After overnight incubation, the two-cell stage embryos were selected and transferred into the oviduct of pseudopregnant ICR females. Pups were genotyped by PCR with two primers (5’-CTCACTATTGAGGTGGTTAGAGGT-3’, 5’-AAACCAGAGACACTCAATTCCAAG-3’) followed by Sanger sequencing to confirm nucleotide substitutions. The identified mutant founders were further back-crossed with wild-type C57BL/6N mice to generate *Eif2b1^N208Y/+^* (N208Y^HET^) mutant mice. Genotyping was performed by Transnetyx using real-time PCR.

### Animal breeding and study

To assess homozygous phenotype, N208Y^HET^ male and female mice were bred and the pups were monitored daily at Taconic Biosciences, Inc. as a service and Calico Life Sciences, LLC. All dead pups were collected and genotyped, along with the weaned mice, to assess genotype-specific lethality. To study the drug effect, 30 mg/kg or 100 mg/kg 2BAct-medicated diet was prepared as previously described (Wong et al., 2019) and provided to N208Y^HET^ breeding pairs upon breeding setup and continued during the nursing period. The same diet without added compound was used as the control diet. Offspring were genotyped, weaned with 30 mg/kg 2BAct-medicated or control diet, and their body weight was monitored and recorded weekly. To study the phenotype of E18.5 embryos, N208Y^HET^ males and females were fed with a control or 30 mg/kg 2BAct-medicated diet a week prior to the mating and the vaginal plug was checked daily after mating. The day the vaginal plug was detected was considered as embryonic day 0.5 (E0.5). Pregnant female mice were euthanized at E18.5 by CO_2_ and a caesarean section was performed to obtain the embryos. The E18.5 embryos were placed on a warm pad, imaged, and monitored for up to 15 minutes while breathing, movement, and foot pinch response were assessed as described (Kammoun et al., 2018), followed by weight measurement before euthanasia for histopathology. Lean and fat mass of individual mice were measured using EchoMRI-100 (EchoMRI, Houston, TX). To score motor deficit, individual mouse was placed on an open field and monitored the presence of abnormal gait, tremor, and other neurological features, then scored its disease severity according to the published clinical scoring system (Traka, 2019). All animal experiments and methods were approved by the Institutional Animal Care and Use Committee of Calico Life Sciences, LLC. and Taconic Biosciences, Inc.

### ELISA

3-4 month old mice were euthanized with CO_2_ and whole blood was collected via cardiac puncture and placed in K2 EDTA-treated BD microtainer tubes. Plasma was collected after spinning at 7000 g for 4 min at 4 ℃ and plasma GDF15 and FGF21 levels were quantified using the GDF15 ELISA (R&D #MGD150) and FGF21 ELISA (BioVender #RD291108200R), respectively, following manufacturer’s instructions. Plasma 2BAct drug exposure was measured as previously described (Wong et al., 2019).

### Embryo or tissue collection

For RNA extraction and western blot analysis, 3-4 month old mice were euthanized with CO_2_, and cerebellum, spinal cord, liver, kidney, lung, muscle and spleen were freshly collected and snap frozen in liquid nitrogen and stored at -80 ℃. For histology and immunohistochemistry, 11-month-old mice were deeply anesthetized and perfused with normal saline followed by 10% buffered formalin via transcardiac perfusion. Brains and spines were excised and post-fixed in 10% buffered formalin for 24-48 hours. To assess the effect of 2BAct removal, 3-month-old female mice were euthanized with CO_2_ and spines were excised and fixed in 10% buffered formalin for 24 hours. For E18.5 embryo histopathology, embryos were fixed with 10% buffered formalin for 24 hours.

### Histology, histopathology, and immunohistochemistry

After fixation, E18.5 embryos were sliced along the transverse plane. Brains were coronally sliced and spinal cords were collected via laminectomy on the spinal column and separated into cervical, thoracic & lumbar regions. Sliced samples were processed for paraffin embedding, sectioned at 5 -6 µm, and mounted on adhesive-coated slides for staining. Histopathological evaluation of E18.5 embryos was carried out by Comparative Pathology Laboratory, University of California Davis School of Veterinary Medicine (Davis, CA, USA) as a service.

Immunohistochemistry was carried out in-house using Leica Bond RX Automated IHC Platform using modified IHC protocol F or by Histobridge, LLC. using Dako Link48 plus as a service with the primary antibodies described in the Key Resources Table and species-appropriate ImmPRESS Polymer Detection Kit (Vector Laboratories) and developed with 3,3’-Diaminobenzidine (DAB) or BOND IHC Polymer Refine Detection (DS9800, Leica Biosystems) followed by hematoxylin counterstaining. Luxol Fast Blue (LFB) was carried out following the manufacturer’s instructions. After staining, sections were dehydrated through successive ethanol solutions, cleared in xylene, and coverslipped using xylene-based mounting media. To examine histopathological timepoints in the brain, sections from two coronal levels of the corpus callosum were examined. For the spinal cord, coronal sections from cervical and thoracic levels were examined. Image capture was achieved using Olympus VS200 slide scanner (Olympus) or NanoZoomer HT2.0. Image analysis was performed using HALO image analysis software (Indica Labs), VS200 Desktop (Olympus), or a QuPath (Bankhead et al., 2017). The same parameters for microscopy and image analysis were uniformly applied to all images for each timepoint and histological staining. For the spinal cord and LFB and MBP staining of corpus callosum, the mean area fraction from two sections from both the cervical and thoracic levels served as the value for each subject to normalize to the different size of the anatomical levels. For other staining of the corpus callosum, positive staining from a fixed area in the corpus callosum of a single section at each coronal level served as the value for each subject.

### Bioanalysis of 2BAct

Plasma, liver, lung, kidney, and brain samples were sent to Quantitative, Translational & ADME Sciences group at Abbvie Lake County for bioanalysis. Samples and standards were extracted by protein precipitation and analytes were separated using C18 reverse phase chromatography prior to analysis with tandem mass spectrometry. Sample concentrations were calculated using the equation derived from regression analysis of the peak area ratio (analyte/internal standard) of the spiked standards versus concentrations.

### RNA extraction

WT, E198K or N208Y MIN-6 cells were plated in 12-well plates in complete media containing 200 nM ISRIB and grown until 80% confluent. ISRIB-containing media was removed, cells were washed once with PBS before media lacking ISRIB was replaced to the wells. ISRIB was withdrawn for 24 hours prior to cell lysis in RT lysis buffer (Qiagen). RNA extraction using the RNeasy Mini Kit (Qiagen) was performed following the manufacturer’s protocol.

Following the indicated animal treatments, the cerebellum, spinal cord, liver, kidney, lung, muscle and spleen were flash frozen in liquid nitrogen prior to RNA extraction. Tissues were treated with RNA-later ICE (Invitrogen #AM7030) for 24 hr at -20C and RNA extraction was performed following the MaxMAX total RNA isolation kit (Invitrogen #AM1830). Briefly, tissues were homogenized in lysis buffer + β-mercaptoethanol using a Qiagen TissueLyser II (Qiagen, Hilden, Germany) for 2 x 2 minute intervals at 25 Hz with the addition of one 7 mm stainless steel bead (Qiagen #69990). 200 uL of tissue lysate was used for KingFisher Flex (Thermo Fisher Scientific, Waltham, USA) RNA extraction according to manufacturer’s protocol, eluting in 60 uL nuclease-free water.

### qRT-PCR

cDNA was synthesized from equal amounts of RNA using the High-capacity cDNA Reverse Transcription Kit (Applied Biosystems #4368814) according to the manufacturer’s protocol. TaqMan qRT-PCR reactions were performed on Quantstudio 6 Flex (Thermo Fisher Scientific, Waltham, MA, USA) using TaqMan Universal PCR Master Mix (Applied Biosystems #4304437) with the following TaqMan Gene Expression Assays: *Atf3* (Mm00476032_m1), *Chop* (Mm01135937_g1), and *Actin* (Mm00607939_s1) purchased from Applied Biosystems. Gene expression for each sample was measured in triplicate. Fold change of gene expression was performed using the comparative CT (-ΔΔ CT).

### Gene expression analysis with nCounter

Multiplex transcript expression levels were measured by nCounter (Nanostring Technologies, Seattle, WA) with a custom panel containing our ISR CLIC genes. 100 - 500 ng of purified total RNA was used for nCounter gene expression analysis as instructed by the manufacturer.

Briefly, reporter and capture probes to the genes of interest were hybridized to total RNA at 65°C for 16 hr. Hybridized probes were then captured to the nCounter cartridge prior to imaging and quantification.

### nCounter data analysis

Raw counts were background subtracted against negative control probes then normalized against housekeeping genes (*B2m*, *Gapdh*, *Hprt*, *Rpl19*) and positive control samples. The counts were log_2_-transformed and a Z-score for each gene was calculated by subtracting the overall mean of the control group from the sample and dividing that result by the SD of all of the measured intensities. ISR pathway activation was measured by taking the averages of the Z-score within each sample. GraphPad Prism (La Jolla, CA) was used to perform statistical analyses utilizing one way ANOVA with Dunnett’s Test performed post-hoc to correct for multiple comparisons. Log_2_ fold changes of ISR CLIC genes were clustered by correlation-based distance using the heatmap.2 function from the gplots package (R Core Team, 2018; Warnes et al., 2005)

### Bulk RNA-seq and analysis

RNA-seq libraries were prepared using purified RNA isolated as described above. RNA quality and concentration were assayed using the 5300 Fragment Analyzer System (Agilent Technologies, Santa Clara, CA, USA). RNA-seq libraries were prepared using the NEBNext Ultra II Directional RNA Library Prep Kit for Illumina (New England Biolabs, Ipswich, MA, USA). Libraries were sequenced on an Illumina NovaSeq 6000 sequencer.

RNA-seq library mapping and estimation of expression levels were computed as follows. Reads were mapped using salmon (Patro et al., 2017), version 1.9.0, to the transcriptome index derived from the mm10 reference mouse genome and the Encode vM12 primary assembly annotation (Mudge and Harrow, 2015).

In order to test for differential expression, for each of the three comparisons, we used sleuth with a design matrix corresponding to two groups (Pimentel et al., 2017). Gene set enrichment was done using fgsea using genes ranked in descending order according to the p value and fold change directionality, as reported by sleuth (Korotkevich et al., 2019).

### GEF assay

WT and MIN-6 N208Y cells were grown to approximately 80% confluence in complete culture media containing 200 nM ISRIB before replacing media containing no drug for 1 hour prior to cell harvest. Cell pellets were washed in PBS, flash frozen in liquid nitrogen and stored at -80C until cell lysis. Each cell pellet was resuspended in 20 mM Hepes, 150 mM KCl, 2 mM TCEP, pH 7.4 + cOmplete EDTA-free protease inhibitor cocktail (Roche). Cells were lysed by bead-based homogenization using Bullet blender (Storm Pro, Next Advance) for 2 x 30 seconds (Setting 10) with 0.15mm zirconium oxide beads. Lysates were clarified by centrifugation at 21,000 x g, 15 min, 4C to remove cellular debris. Protein concentration was determined by BCA and lysates were aliquoted and stored at -80C until use. Bodipy-FL-GDP loaded eIF2 was used as a substrate for in vitro GEF assay. 1 mg/mL protein lysates (unless otherwise described) were treated in vitro with indicated ISRIB or 2BAct concentrations, 25 nM Bodipy-FL-GDP loaded eIF2, 0.1 mM GDP, and 1 mg/mL BSA. Fluorescence decay was measured on a SpectraMax i3x plate reader (Molecular Devices) with the following parameters: plate temperature = 25C; excitation wavelength = 485 nm (15 nm width); emission wavelength = 535 nm (25 nm width); read duration = 30 min at 45 s intervals. Data were analyzed in Prism fitting to a single exponential decay curve to calculate GDP half-lives.

## Acknowledgements

We thank Karla Barron, Samira Nazertehrani, Paulyn Cha, Ellie Karlsson, Wendy Craft, Baby Martin-McNulty, Ikenna Anigbogu, Andrew Keyser, Amy Jo Johnson, David Hendrickson, Arthur N. Nikkel, and the Non-regulated Bioanalytical team at AbbVie, Lake County, IL for experimental support. This work was supported by Calico Life Sciences LLC.

