## Supplementary Materials for "eIF2B Activator Rescues Neonatal Lethality of an eIF2Bα Sugar Phosphate Binding Mutation Associated with Vanishing White Matter Disease"

### Supplementary Material

**Figure Supplement 1.**
(A) Immunoblot of purified recombinant eIF2Bα WT, E198K or N208Y against the eIF2Bα antibody.

**Figure Supplement 2.**
(A) Number of P0 pups found dead or alive in each genotype as shown in Figure 2C.
(B) Foot pinch responsiveness of E18.5 embryos for each genotype as shown in Figure 2E.
(C) Representative H & E images of pancreas, spleen, thymus, and brown fat. scale bars, 50 µm.

**Figure Supplement 3.**
(A) Body weight measurements of WT, N208Y$^{HOM}$, and N208Y$^{HET}$ mice with 2BAct. Male WT, n = 6; N208Y$^{HET}$, n = 14, WT + 2BAct, n = 2, N208Y$^{HET}$ + 2BAct, n = 9; N208Y$^{HOM}$ + 2BAct, n = 4.
(B) Body weight measurements of 4-week-old male WT and N208Y$^{HOM}$ mice with 2BAct. WT + 2BAct, n = 44; N208Y$^{HOM}$ + 2BAct, n = 32.
(C) Fat and lean mass measurement of 3-month-old female mice by EchoMRI. WT + control, n = 8; WT + 2BAct, n = 13; N208Y$^{HOM}$ + 2BAct, n = 7.
Male daily food consumption (D) and the percentage of daily food consumption per body weight (E) measured between 5 and 6 weeks of age.WT + 2BAct, n = 44; N208Y$^{HOM}$ + 2BAct, n = 32.
(B) to (E) Error bars are Standard deviation. Student's t-test. **** p < 0.0001, ns = not significant.
Representative H & E, OLIG2, IBA1, GFAP, and ATF3 IHC images and quantification of the thoracic region of the spinal cord (F) and the corpus callosum region of the brain (G). Scale bars, 500 µm. Inset scale bars, 50 µm. (B and C) WT+Ctrl, n = 6 (3 females and 3 males); WT+2BAct, n = 4 (2 females and 2 males); N208Y$^{HOM}$ + 2BAct, n = 4 (2 females and 2 males). Error bars are Standard deviation. One-way ANOVA with Holm-Sidak's multiple comparisons test. * p < 0.05, ** p < 0.01, *** p < 0.001, **** p < 0.0001, ns = not significant.

**Figure Supplement 4.**
(A) Average z-core of the ISR CLIC genes calculated from nCounter gene expression profiling of various tissues from 4-month-old male mice WT + 2BAct-medicated diet normalized to WT + control det. WT + control diet, n = 3 and WT + 2BAct-medicated diet, n=3. Error bars are Standard deviation. Two-way ANOVA with Holm-Sidak's multiple comparisons test. ns, not significant.
(B) Immunoblot of eIF2B subunits using lung (left) and kidney (right) lysates of 2BAct-treated WT and N208Y$^{HOM}$ mice.
(C) Quantification of bands in (B) normalized to eIF2α expression and represented as % of WT expression. n=4. Error bars are Standard deviation. Two-way ANOVA with Holm-Sidak's multiple comparisons test. *** p < 0.001, **** p < 0.0001.
(D) 2BAct $EC_{50}$ in MIN6 N208Y cell line was calculated from measuring ATF4 suppression from a 4 hr dose response of 2BAct following a 24 hr withdrawal of ISRIB. Immunoblot of ATF4 protein was normalized to eIF2α (left) and displayed as relative abundance (right), n = 3.

**Figure Supplement 5.**
(A) Percentage of body weight change of 3-month-old female mice after 2BAct withdrawal. WT + 2BAct, n = 6; WT + 2BAct withdrawal, n = 7; N208Y$^{HOM}$ + 2BAct, n = 3; N208Y$^{HOM}$ + 2BAct withdrawal, n = 4

**Figure Supplement 7.**
(A) Volcano plot demonstrating gene expression changes between 2BAct and CHOW in WT cerebellum. Red dots show ISR CLIC genes and the dotted line indicates significance threshold (adj $p < 0.05$).

**Supplemental Table 1.**
Fold-change of ISR CLIC gene expression panel for 2BAct-treated N208Y$^{HOM}$ compared to WT 2BAct-treated in cerebellum and spinal cord. Two-way ANOVA with Holm-Sidak's multiple comparison test. * $p < 0.05$, ** $p < 0.01$, *** $p < 0.001$, **** $p < 0.0001$

**Supplemental Table 2.**
Fold-change of ISR CLIC gene expression panel for N208Y$^{HOM}$ 2BAct withdrawal compared to WT 2BAct-treated in cerebellum, spinal cord, kidney, lung, muscle, liver, and spleen. n=4. Two-way ANOVA with Holm-Sidak's multiple comparison test. * $p < 0.05$, ** $p < 0.01$, *** $p < 0.001$, **** $p < 0.0001$

**Supplemental Table 3.**
List of genes differentially expressed in the comparison of R191H$^{HOM}$ vs WT and N208Y$^{HOM}$ + 2BAct vs WT + 2BAct, as shown in Figure 7D.

**Supplemental Table 4.**
List of gene sets significantly over-represented in all three comparisons as shown in Figure 7E or only in N208Y$^{HOM}$ 2BAct withdrawal vs WT + 2BAct comparison as shown in Figure 7F. The gene sets with values of Normalized Enrichment Score (NES) > 2.5 and adjusted $p < 0.05$ were shown.

A

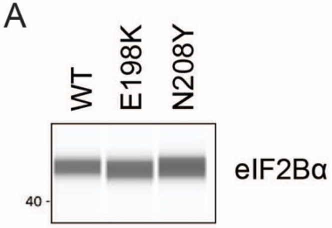


Figure S1

A

**P0 pups viability**

| | WT | N208Y$^{HET}$ | N208Y$^{HOM}$ |
|---|---|---|---|
| Live | 21 (100 %) | 44 (95.7 %) | 2 (11.1 %) |
| Dead | 0 ( 0 %) | 2 ( 4.3 %) | 16 (88.9 %) |

B

**E18.5 embryo viability**

| Pinch response | WT | N208Y$^{HET}$ | N208Y$^{HOM}$ |
|---|---|---|---|
| Yes | 13 (100 %) | 30 (96.8 %) | 13 (86.7 %) |
| No | 0 ( 0 %) | 1 ( 3.2 %) | 2 (13.3 %) |

C

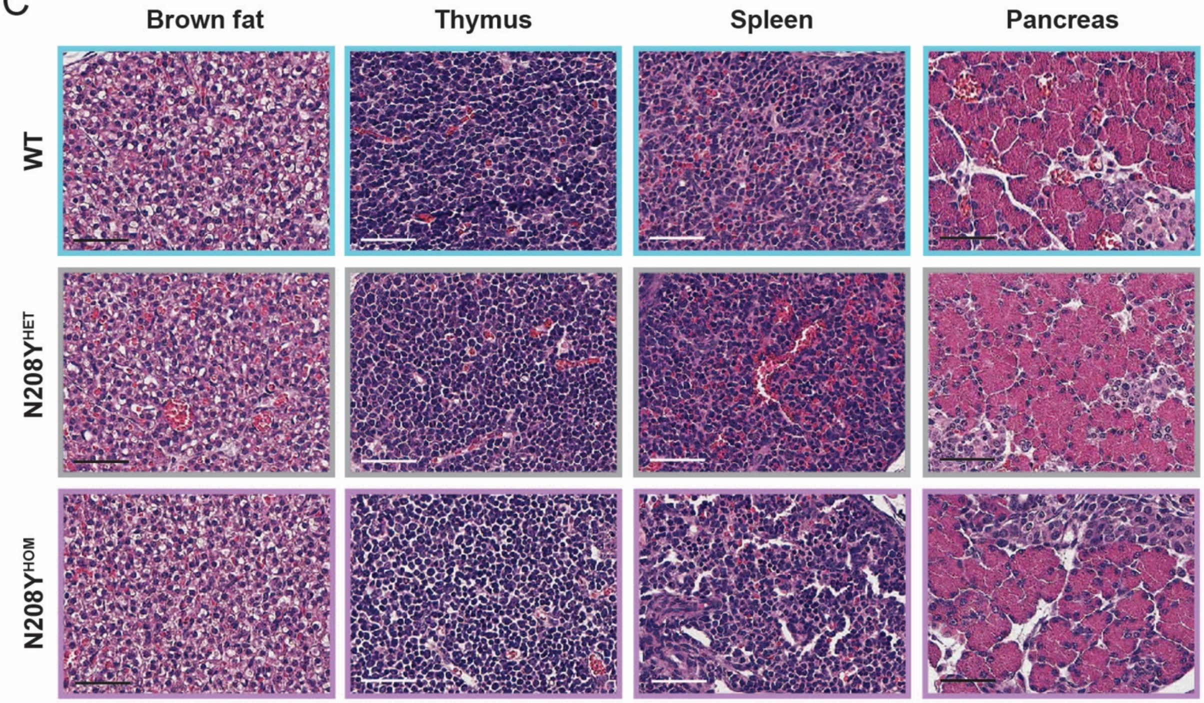


### Figure S2

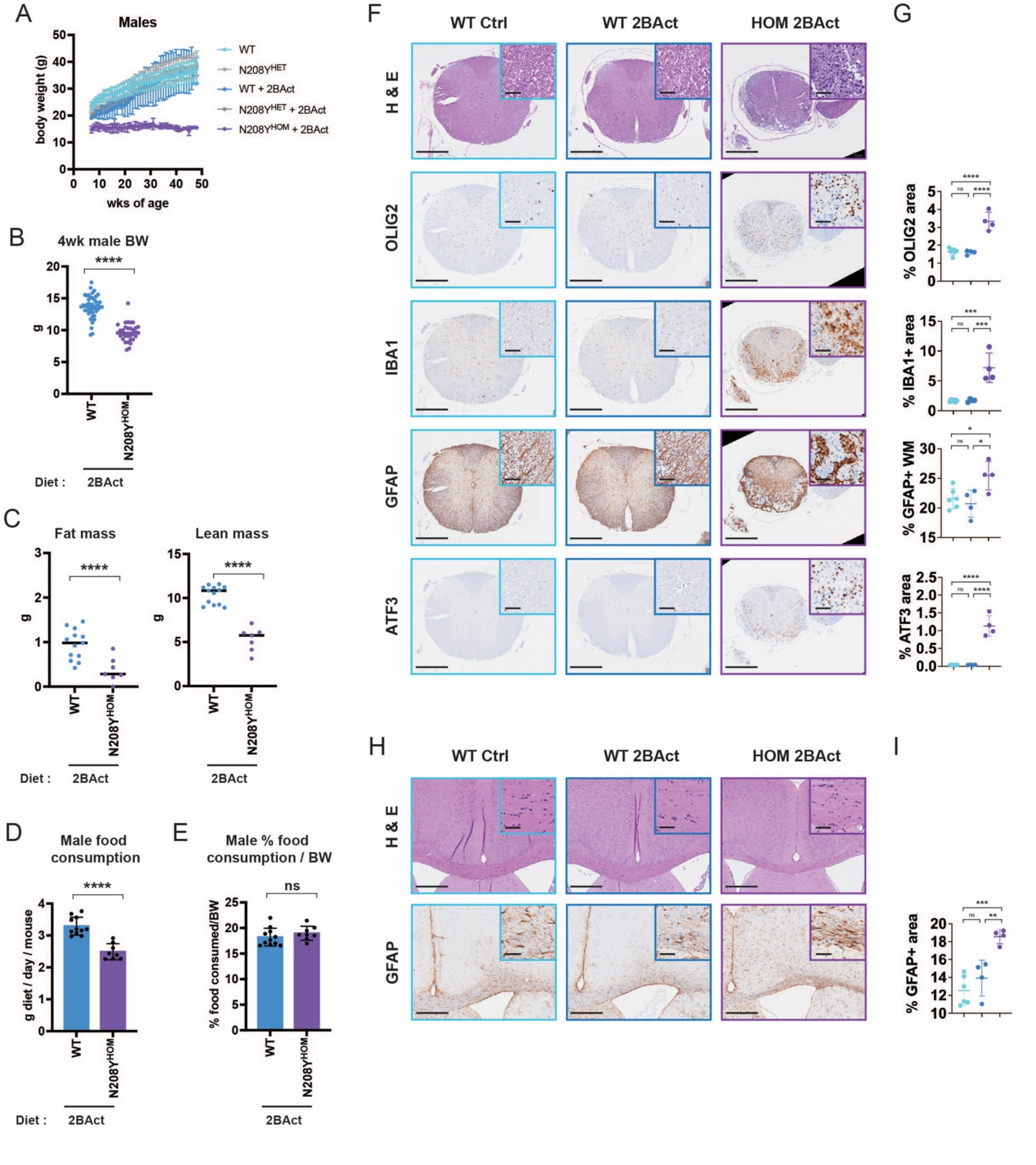
A
Males
body weight (g)
wks of age
WT
N208Y^HET
WT + 2BAct
N208Y^HET + 2BAct
N208Y^HOM + 2BAct
B
4wk male BW
Diet : 2BAct
C
Fat mass
Lean mass
D
Male food consumption
g diet / day / mouse
E
Male % food consumption / BW
% food consumed/BW
F
WT Ctrl
WT 2BAct
HOM 2BAct
H & E
OLIG2
IBA1
GFAP
ATF3
G
% OLIG2 area
% IBA1+ area
% GFAP+ WM
% ATF3 area
H
I
% GFAP+ area

### Figure S3

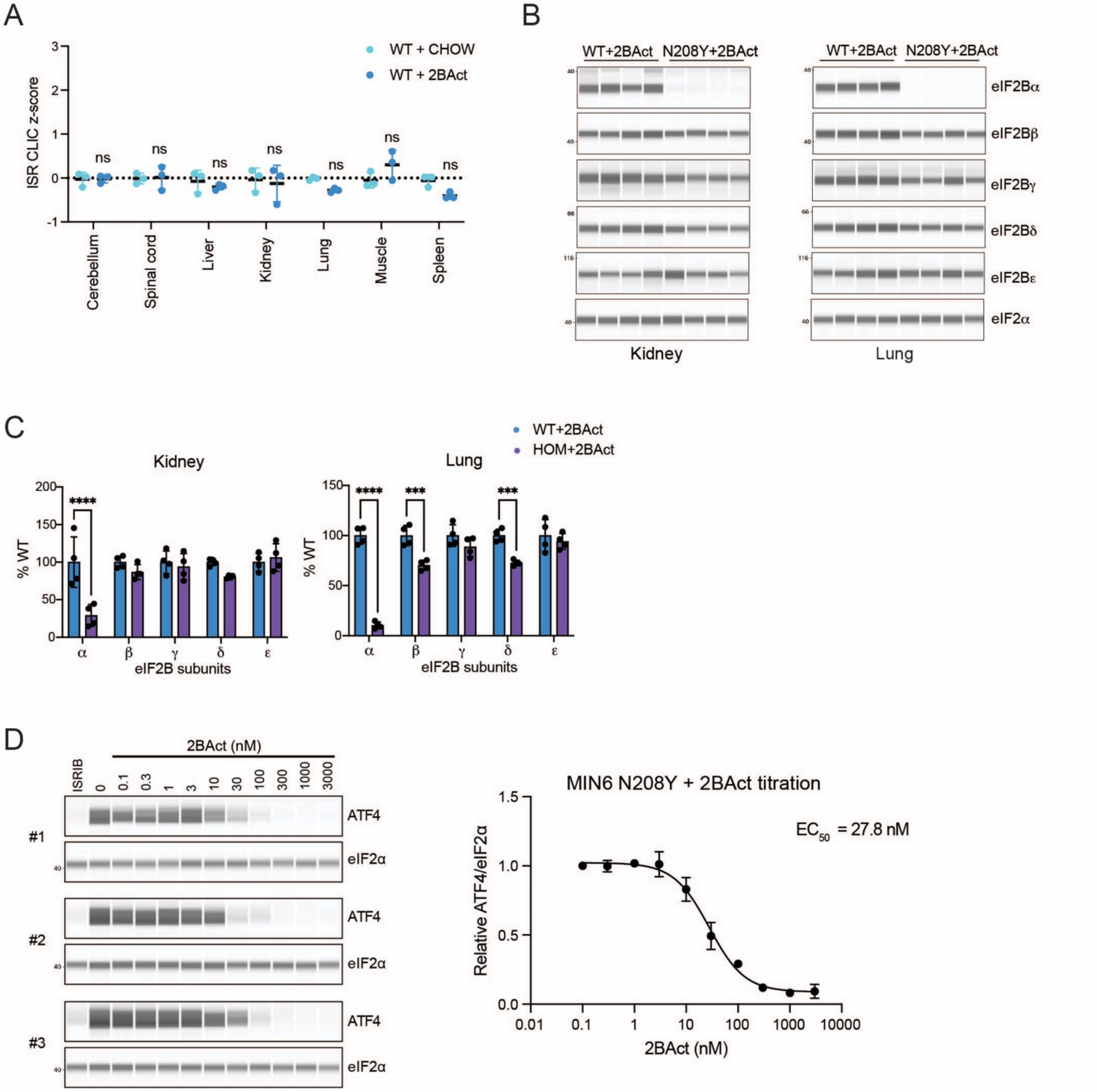

A
WT + CHOW
WT + 2BAct
ISR CLIC z-score
ns
Cerebellum
Spinal cord
Liver
Kidney
Lung
Muscle
Spleen
B
WT+2BAct
N208Y+2BAct
eIF2Bα
eIF2Bβ
eIF2Bγ
eIF2Bδ
eIF2Bε
eIF2α
Kidney
Lung
C
WT+2BAct
HOM+2BAct
Kidney
Lung
% WT
eIF2B subunits
D
2BAct (nM)
ISRIB
ATF4
eIF2α
#1
#2
#3
MIN6 N208Y + 2BAct titration
EC50 = 27.8 nM
Relative ATF4/eIF2α
2BAct (nM)


### Figure S4

A

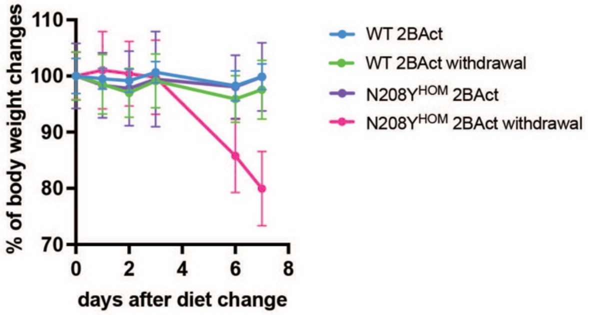
% of body weight changes
110
100
90
80
70
0
2
4
6
8
days after diet change
WT 2BAct
WT 2BAct withdrawal
N208Y$^{HOM}$ 2BAct
N208Y$^{HOM}$ 2BAct withdrawal

### Figure S5

A

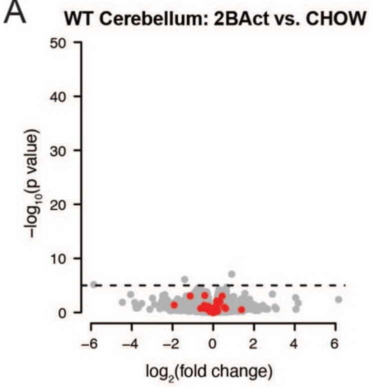
WT Cerebellum: 2BAct vs. CHOW
50
40
30
20
10
0
−log10(p value)
−6
−4
−2
0
2
4
6
log2(fold change)

### Figure S7
